# The guide RNA sequence dictates the slicing kinetics and conformational dynamics of the Argonaute silencing complex

**DOI:** 10.1101/2023.10.15.562437

**Authors:** Peter Y. Wang, David P. Bartel

**Affiliations:** Whitehead Institute for Biomedical Research, 455 Main Street, Cambridge, MA, 02142, USA; Howard Hughes Medical Institute, Cambridge, MA, 02142, USA; Department of Biology, Massachusetts Institute of Technology, Cambridge, MA, 02139, USA

**Keywords:** RNAi, Argonaute, AGO2, RISC, slicing, microRNA, siRNA, kinetic analysis, RNA–protein interactions

## Abstract

**SUMMARY:** The RNA-induced silencing complex (RISC), which powers RNA interference (RNAi), consists of a guide RNA and an Argonaute protein that slices target RNAs complementary to the guide. We find that for different guide-RNA sequences, slicing rates of perfectly complementary, bound targets can be surprisingly different (>250-fold range), and that faster slicing confers better knockdown in cells. Nucleotide sequence identities at guide-RNA positions 7, 10, and 17 underlie much of this variation in slicing rates. Analysis of one of these determinants implicates a structural distortion at guide nucleotides 6–7 in promoting slicing. Moreover, slicing directed by different guide sequences has an unanticipated, 600-fold range in 3′-mismatch tolerance, attributable to guides with weak (AU-rich) central pairing requiring extensive 3′ complementarity (pairing beyond position 16) to more fully populate the slicing-competent conformation. Together, our analyses identify sequence determinants of RISC activity and provide biochemical and conformational rationale for their action.

**HIGHLIGHTS:**

- Sequence of guide RNA can alter slicing rate of fully paired substrate by 250-fold
- Sequences that cause more rapid slicing direct more efficient RNAi in cells
- Strong central pairing imparts tolerance for mismatches to the guide 3′ region
- This tolerance is attributable to more fully populating the slicing conformation

## INTRODUCTION

Metazoan Argonaute (AGO) proteins associate with guide RNAs and downregulate transcripts that pair to the guides, which can be either microRNAs (miRNAs) or small interfering RNAs (siRNAs). Pairing that is less extensive, often involving recognition of only the seed region (guide-RNA nucleotides 2–8), leads to recruitment of TNRC6, which in turn recruits deadenylation complexes, ultimately causing degradation and/or translational repression of the target,^1,2^ whereas extensive pairing can also lead to AGO-catalyzed endonucleolytic slicing of the target transcript.^3–5^

Slicing is the biochemical basis of RNA interference (RNAi), which provides defense against viruses and transposable elements for many eukaryotic species.^6,7^ In this context, a slicing-competent AGO, together with its guide RNA, is called an RNA-induced silencing complex (RISC).^8,9^ Of the four mammalian AGO paralogs, AGO2 is the most adept at catalyzing slicing.^10–12^ In mammals, 20 miRNAs direct AGO2-catalyzed slicing of 24 transcripts,^13–16^ and in mice, endogenous siRNAs also direct slicing required for oocyte meiotic maturation.^17^ Slicing is also required for the biogenesis and maturation of at least two miRNAs^18–21^ and most siRNAs.^22–24^ Moreover, slicing is key to potent knockdown directed by synthetic siRNAs, which are useful research reagents and increasingly applied to treat disease, with six siRNA therapies now FDA approved.^25^

The fully paired, slicing conformation was initially proposed to occur through a two-step process, in which pairing begins at a pre-organized guide-RNA seed region and then propagates contiguously, like a zipper, until reaching the guide 3′ terminus.^26–^ _^32^_ However, structural studies of eukaryotic AGO proteins show that only part of the seed region is pre-organized in a conformation suitable to nucleate pairing.^33–35^ Moreover, simple propagation of pairing from the seed is hindered by a narrowing of the AGO central cleft.^36^ Furthermore, contiguous propagation of pairing from a stationary seed to form ∼20 pairs, in which the guide RNA has wrapped nearly twice around the target, poses topological challenges, when considering that the unpaired guide contacts AGO throughout its length.^30^ To address these issues, the model has been revised to be a four-step process (Figure 1A).^2^ In the first step, pairing nucleates at pre-organized nucleotides 2–5, and in the second step, it propagates to the remainder of the seed region.^36–40^ In step three, instead of pairing continuing to zipper from the seed through the remainder of the guide, another nucleation event creates a second guide−target helix, located at the 3′ supplementary region of the guide, which tends to center at positions 13−16.^2^ In step four, this second helix rotates as additional pairing forms on both of its flanks (Figures 1A and S1A).^2^ Propagation of pairing from the second helix to the end of the guide helps to disrupt remaining contacts between the 3′ region of the guide and AGO, including contacts between the 3′ terminus and the PAZ domain, thereby untethering the second duplex, allowing it to rotate and migrate over the N domain and into the AGO central cleft. This rotation and movement of the second helix enables pairing to form between the two helices (propagating from either or both directions), ultimately completing a continuous double helix (Figures 1A and S1A).^2^ Recent structures support this four-step model (Video S1).^41–43^

**Figure 1.**
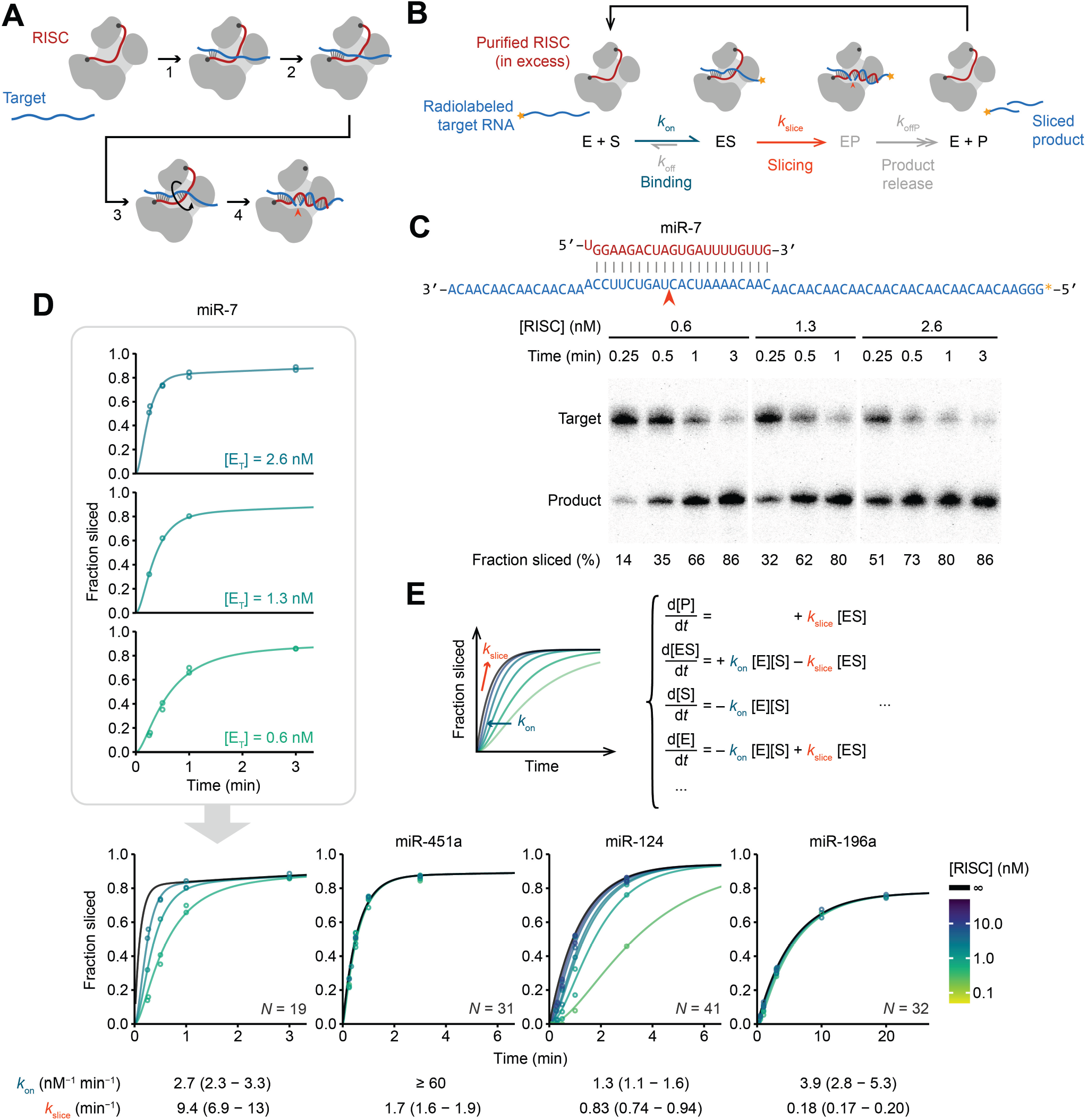
Precise measurement of RISC-catalyzed slicing kinetics. (A) Schematic of the four-step model for RISC engagement with fully complementary targets. See text for description. (B) Minimal kinetic scheme of binding (teal) and slicing (orange). Colors are as in (A). E represents RISC; S represents target; P represents sliced product, and the orange caret represents the AGO2 active site. For single-turnover reactions, RISC is at high concentrations, in large excess over target, such that product release and substrate dissociation (both gray) have negligible impact. (C) Representative results of miR-7-directed slicing of a perfectly complementary target. Site of slicing is indicated (orange caret); colors otherwise as in (A). Initial target concentration was 0.05 nM. (D) Results of slicing assays evaluating four different miRNAs. Data points and reaction curves from ODE model fitting are plotted in a gradient from green to purple for increasing RISC concentrations. The extrapolated reaction curve at infinite RISC concentration, which represents a reaction rate determined purely by slicing kinetics, is in black. Time points beyond the limits of the x-axes are not shown. *N* indicates the number of data points measured for each miRNA. Values of *k*_on_ and *k*__slice__ obtained from ODE model fitting are shown for each miRNA. Ranges in parentheses indicate 95% CIs from model-fitting. Binding rate for miR-451a was too fast to resolve in our assays and was thus set to the diffusion limit (60 nM^−1^ min^−1^). All miRNAs were 22-nt long, unless stated otherwise. (E) Simplified ODE model describing the reaction in (B), with a schematic kinetic plot demonstrating the contribution from each kinetic parameter. Colors are as in (B) and (D). See also Figures S1, S2, S3, and S4, Table S1, and Video S1.

RISC biochemical studies have measured kinetic parameters and defined base-pairing requirements of slicing guided by either let-7a or miR-21.^39,44–47^ However, the extent to which these findings generalize to other guide-RNA sequences has been unclear. Indeed, less thorough analyses of RISC guided by other RNAs suggest that guide-RNA identity might influence slicing efficiency.^38,48^ Likewise, guide RNAs in high-resolution structure models are limited to only a few sequences, and the impact of different guide-RNA sequences on RISC conformational dynamics has been largely unexplored.

Here, we measured the slicing kinetics of AGO2 loaded with a variety of guide RNAs. We found that the sequence of the guide not only influences binding of the target and products but also dictates the conformation and slicing rate of target-bound RISC, thereby affecting RNAi efficacy in cells.

## RESULTS

### Rate of target association can obscure slicing kinetics

To explore how the guide-RNA sequence influences AGO2-catalyzed slicing, we measured the slicing kinetics of AGO2 loaded with 13 different miRNAs, most of which had been either previously biochemically studied or shown to carry out cellular slicing-mediated functions.^12,13,16,47–49^ Each of the 13 RISCs were generated in cellular extracts, purified by antisense capture and affinity elution,^50^ and quantified using stoichiometric binding of radiolabeled RNA (Figures S1B and S1C). Slicing kinetics of each RISC were then measured using single-turnover slicing assays, with RISC in excess of radiolabeled target RNA harboring a perfectly complementary site for the miRNA (Figure 1B). At each time point, the reaction was quenched and unsliced substrate and sliced product were resolved on denaturing gels (Figure 1C).

The use of single-turnover conditions, with enzyme in excess over substrate, coupled with slow dissociation of fully complementary substrate,^39^ ensured that the reaction kinetics would not be perceptibly influenced by rates of either product release or substrate dissociation and instead would be influenced by rates of substrate association and/or slicing, described by elemental rate constants *k*__on__ and *k*__slice__, respectively (Figure 1B). Previous analyses of AGO2 associated with either let-7a or miR-21 suggested that under typical experimental concentrations of RISC, slicing is slower than association, such that the apparent single-turnover reaction rates correspond to the rate-limiting *k*__slice__ (Figure 1B).^45,47^ However, for some miRNAs that we tested, reaction kinetics increased with RISC concentration, indicating some contribution by second-order association kinetics, i.e., by *k*__on__ (Figure 1D). To separate the contribution of *k*__on__ from that of *k*__slice__, we conducted assays at multiple RISC concentrations and fit the combined results to a simplified ordinary differential equation (ODE) model that included both association and slicing kinetics (Figure 1E).

To validate the results of our ODE model, we used filter binding to separate bound target RNA from free target RNA at different time points (Figures S2A and S2B). At early time points, partial binding of target RNA was observed for miRNAs that had slower association kinetics in our slicing assays but not for those that had faster association kinetics (Figure S2C). These results also confirmed that, as assumed in our modeling, fraction-sliced endpoints falling below 100% could be attributed to a small fraction of defective enzyme (Figures S2D−G). To confirm that the ODE model generated accurate *k*__on__ values, we directly measured association kinetics using filter-binding assays of target RNAs harboring central mismatches, which cannot be sliced. As expected, these *k*__on__ values corresponded to those obtained from ODE modeling of slicing reactions (Figure S3A). Predicted folding of target sequences supported the idea that the *k*__on__ values were largely dictated by structural accessibility of the seed-match region of the target (Figures S3B−D).^38,49,51–54^ Moreover, *k*__on__ values did not correlate with *k*__slice__ values (Figure S3E), which indicated that we had disentangled them from each other to generate accurate *k*__slice__ values for a broad range of miRNAs, enabling investigation of the interplay between miRNA sequence and *k*__slice__ (Figure S4; Table S1).

### miRNA sequence can confer >250-fold differences in *k*_slice_ values of perfectly complementary targets

When presented with their perfectly matched targets, the 13 characterized RISCs had *k*__slice__ values spanning >250-fold, with time of slicing (*τ*__slice__) spanning 6.4 s (95% confidence interval [CI], 4.7–8.7 s) to 28 min (95% CI, 26–29 min) (Figure 2A). Examining sequences of the 13 miRNA guides suggested that increased *k*__slice__ values were potentially associated with a purine (R) at position 10 and an A or U (W) at positions 7 and 17 (Figure 2A).

**Figure 2.**
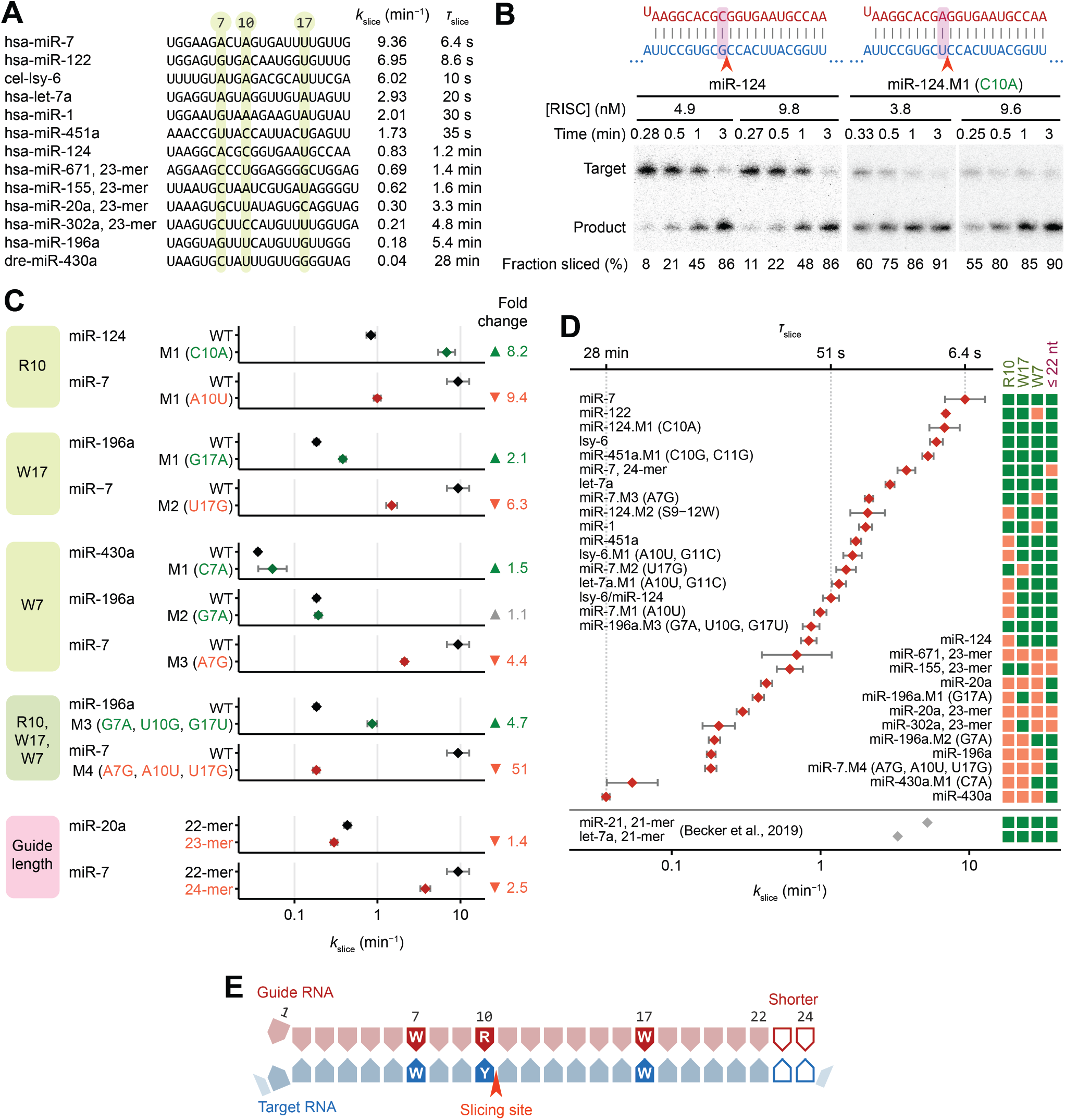
Sequence determinants of slicing rates. (A) A broad range in *k*__slice__ values. The 13 initially tested miRNAs, listed in order of their *k*_slice_ and *τ*_slice_ values. Positions of candidate sequence determinants are highlighted in green. (B) Effect of substituting the base pair at position 10. Shown are representative results for a wildtype miRNA (miR-124) and its mutant (miR-124.M1), in which the C at position 10 was substituted with A (C10A). Initial concentration of target RNA was 0.1 nM in these assays; otherwise, as in Figure 1C. (C) Validation of sequence determinants through base-pair substitutions and additions. Results for wildtype–mutant pairs used to interrogate each sequence determinant are grouped together, with the sequence determinant indicated on the left. *k*_slice_ values are plotted, with error bars indicating 95% Cis from model-fitting. Wildtype values are in black; those for substitutions expected to increase or decrease *k*_slice_ are in green and orange, respectively. Relative fold-change values observed between wildtype and mutants are on the right; a value smaller than the 95% CI of biological replicates is in gray (Figure S5A). (D) Replotting of *k*_slice_ values for perfectly complementary targets for all 29 guides tested in this study. *k*_slice_ values are plotted as in (C), with the top axis also showing the corresponding *τ*_slice_ values, labeling the minimum, maximum, and median values. For reference, *k*_slice_ values from a previous study^47^ are shown at the bottom. On the right, green indicates the presence of a favorable determinant, and orange its absence. (E) Summary of determinants identified for *k*_slice_ of perfectly complementary targets. Colors are as in (B). See also Figure S5 and Table S1.

To test these candidate sequence determinants, we made single-nucleotide substitutions and measured their impact on *k*__slice__ for the corresponding perfectly complementary targets. The purine/pyrimidine identity at position 10 most strongly affected *k*__slice__, conferring an 8.2- or 9.4-fold difference when changing to a purine or vice-versa (Figures 2B and 2C). Changing position 17 to or from an A or U had a 2.1- or 6.3-fold effect on *k*__slice__ (Figure 2C), and changing position 7 from an A to a G reduced *k*__slice__ 4.4-fold, whereas changing it from a G or C to an A increased *k*__slice__, albeit modestly (1.1- or 1.5-fold) (Figure 2C). These changes in *k*__slice__ were generally greater than the variation found in biological replicates (Figure S5A), and triple mutants with concordant substitutions at all three determinants imparted a 4.7- or 51-fold difference in *k*__slice__, further validating the sequence features (Figure 2C). Moreover, most of these effects on *k*__slice__ were stronger than that of a phosphomimetic substitution of serine-387 of AGO2 (Figures S5B and S5C), a residue that, when phosphorylated, is reported to downregulate slicing-mediated repression in cells.^55^

Shorter guide lengths can introduce stronger tension at the miRNA 3′ terminus upon pairing to the miRNA 3′ region.^41^ Reasoning that this tension might favor the release of the 3′-end of the guide, which is a conformational requisite of slicing,^32^ we measured *k*__slice__ values of different length isoforms for two guides and found small but consistent increases in *k*__slice__ for shorter guides (Figure 2C).

Together, these results supported the identification of four determinants that influence the slicing step of perfectly complementary target RNAs. They were, in decreasing order of effect size: (1) a purine at miRNA position 10, (2) an A or U at miRNA position 17, (3) an A or U at miRNA position 7, and (4) a shorter miRNA (Figures 2C−E). Moreover, when compared to the 2-fold range in *k*__slice__ values reported previously for perfectly complementary sites of different guide RNAs,^47^ the >250-fold range in *k*__slice__ values observed here indicated that these values can span a much larger range and are often slower than previously appreciated (Figure 2D).

### *k*_slice_ can limit multiple-turnover catalysis for fully complementary targets

Studies of RISCs programmed with several different guide RNAs indicate that for perfectly complementary targets, the rate of product release (*k*__offP__) limits the rate of multiple-turnover catalysis,^9,44,46,48^ but with introduction of multiple mismatches, slicing can become rate limiting.^39,44^ The wide range of *k*__slice__ values observed when testing additional miRNAs (Figure 2A) suggested that, for many guides, mismatches might not be required for slicing to limit the rate of multiple-turnover catalysis. To test this possibility, we conducted multiple-turnover slicing assays for several guides to measure *k*__offP__ values and reaction kinetics. Although some guides, such as miR-7.M1, had burst-phase kinetics indicating *k*__offP__-limited catalysis, others, such as miR-196a.M3, had single-phase kinetics limited by *k*__slice__ throughout multiple rounds of catalysis, which indicated that *k*__offP__ exceeded *k*__slice__ (Figures S5D and S5E). These results showed that, depending on the miRNA sequence, either slicing or product release can dictate the maximal catalytic rate constant for perfectly complementary targets of AGO2 RISC.

### Determinants that enhance *k*_slice_ promote greater knockdown efficacy in cells

In-cell screens have identified siRNA sequence features that confer greater knockdown efficacy, which provide guidelines for siRNA design^56^ (Figure S5F). Although some of these features are explained by either more efficient maturation of RISC^57,58^ or increased structural accessibility of target sites,^59^ three features (non-G7, A10, U17) have remained mechanistically unexplained. These three features aligned precisely with three of our *k*__slice__ determinants (W7, R10, W17) (Figures 2E and S5F), which implied that more rapid AGO2-catalyzed slicing underlies the functional efficacy of siRNAs bearing these features.

To validate the functional impact of *k*_slice_ sequence determinants on RNAi efficacy, we used a dual-luciferase reporter assay (Figure 3A). We tested two pairs of wildtype and mutant miRNAs, in which mutants had concordant single-nucleotide substitutions at positions 7, 10, and 17 that either reduced or increased *k*_slice_ (Figure 3B). For each guide tested, we placed the corresponding perfectly complementary site within two different open-reading-frame (ORF) contexts (either 3xFLAG−sfGFP or 3xHA−human-orthogonal SUMO). Translation rates of these ORFs were placed under control of an iron-response element (IRE), such that the translation initiation rate could be tuned by addition of either iron or an iron chelator^60^ (Figures 3A and 3C), which enabled testing of whether kinetic competition by translocating ribosomes expected to clear bound RISCs might contribute to the benefit of faster slicing.^61^ Effects on reporter RNA levels were monitored using destabilized NanoLuc luciferase translated from a downstream ORF under translational control of an EMCV internal ribosome entry site (IRES), such that NanoLuc activity reported on the levels of the reporter transcript without sensitivity to IRE modulation of cap-dependent translation initiation (Figure 3A).

**Figure 3.**
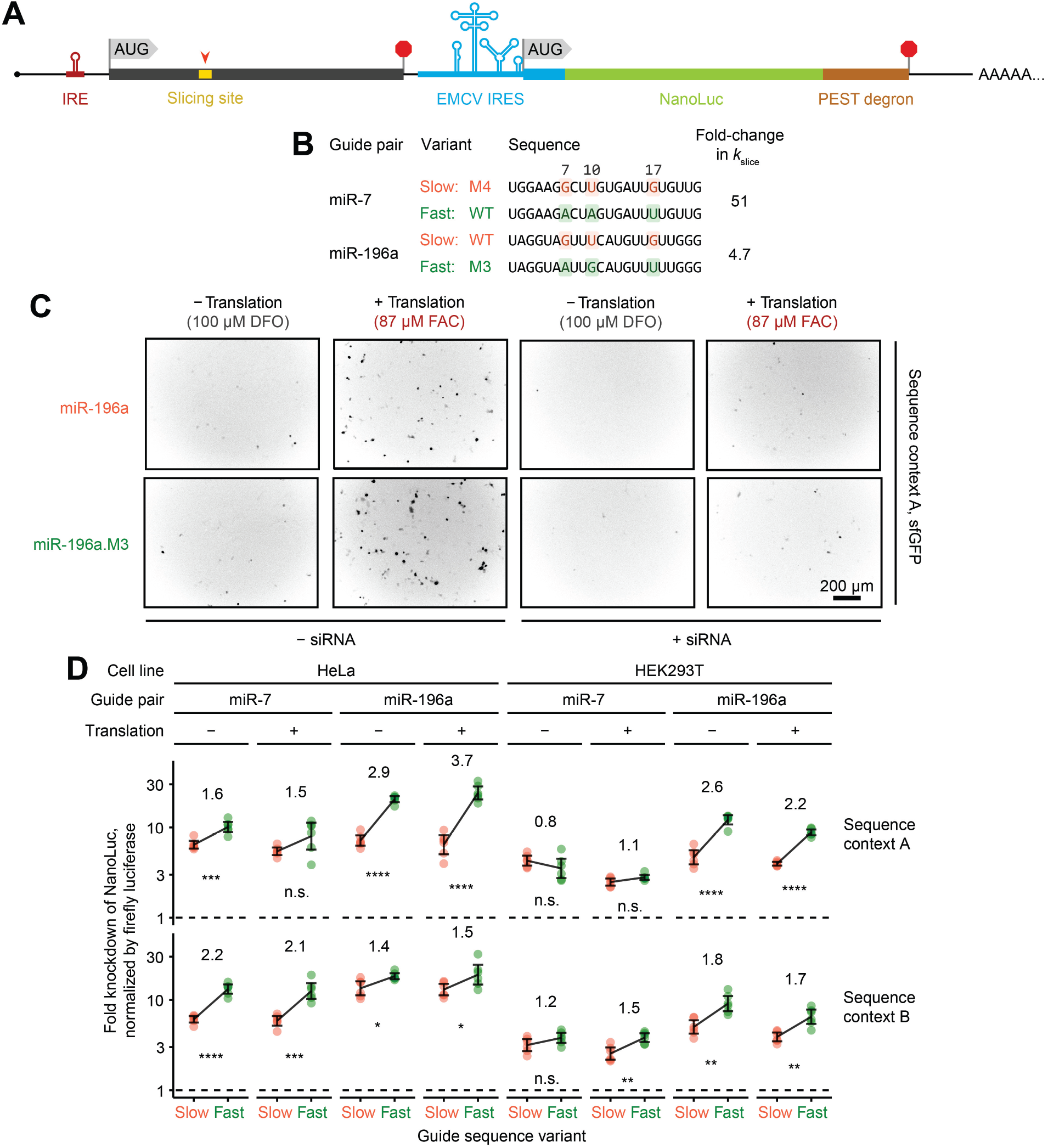
Favorable sequence determinants confer stronger knockdown in cells. (A) Schematic of reporter constructs. Thicker bars indicate ORFs; thinner bars indicate RNA elements; AUGs indicate start codons; stop signs indicate stop codons, and orange caret points to the perfectly complementary target site in yellow. The target site resided in one of two different ORF contexts—either context A, in which it was positioned between a region coding for the 3xFLAG-tag and a region coding for superfolder GFP (sfGFP),^78^ or context B, in which it was positioned between a region coding for the 3xHA-tag and a region coding for human-orthogonal SUMO protein (SUMO^Eu1^).^79^ (B) Sequences of four guides tested with reporter assays, shown as two pairs of wildtype and mutant sequences. Key features of the slower-slicing variant are in orange, and those of the faster-slicing variant in green. (C) Representative fluorescent microscopy images of HEK293T cells transfected with reporter constructs containing ORF context A, which included sfGFP. Cells were imaged after 17 h of treatment with either iron chelation (desferroxamine; DFO) or iron (ferric ammonium citrate; FAC). Scale bar is in the bottom right; all images were acquired at the same magnification. Colors were inverted for visibility. (D) Greater knockdown by guides with more favorable slicing determinants. Plotted are reporter knockdown efficacies observed with the indicated cell lines, treatments, and reporter contexts. Fold differences observed between mean knockdown values of the slow and fast guide variants are shown above each set of data points. Each error bar indicates 95% CIs of the geometric mean across six biological replicates. Significance, as measured using unpaired two-sample *t*-test, calculated in log-scale for fold-change values, is indicated below each set of data points (**** *p* < 0.0001, *** *p* < 0.001, ** *p* < 0.01, * *p* < 0.05, n.s. not significant). Colors are as in (B).

In both HeLa and HEK293T cells, across both reporter sequence contexts, and for both pairs of guides tested, the faster-slicing guide variant usually achieved significantly greater target knockdown (Figure 3D). The smallest differences were observed for miR-7 in HEK293T cells, which was attributable to higher levels of endogenous miR-7 in these cells,^62^ which might have diminished the effect of transfected miR-7. Interestingly, neither a global change in knockdown efficacy nor an additional advantage of fast-slicing guides was observed when translation through the site of cleavage was substantially increased (Figures 3C and 3D), which suggested that, at least in the context of our reporters, kinetic competition with translocating ribosomes did not affect AGO2-mediated slicing. The minimal effects of translation also suggested that translocating ribosomes did not occlude binding to target sites and did not disrupt secondary structure that was occluding target-site accessibility, although combinations of offsetting effects could not be ruled out. Most importantly, these results established a functional link between *k*_slice_ sequence determinants and RNAi efficacy in cells.

### Formation of continuous helix does not limit *k*_slice_ of fully complementary targets

To examine the mechanism by which the sequence determinants influence *k*_slice_, we probed their effect on the fraction of RISC that achieves the centrally paired, slicing-competent state. Formation of these central pairs and associated conformational changes are thought to be energetically challenging, presumably due to either the protein conformational strain or disruption of favorable contacts between guide and protein that occur upon achieving this slicing-competent configuration (Figure S1A).^41,43^ Thus, *k*_slice_ is proposed to be limited by the fraction of complex populating the centrally paired configuration,^29,63^ as described by *K*_con_, the equilibrium constant defined as the rate constant for forming the centrally paired conformation (*k*_con+_) divided by the rate constant of its reversal (*k*_con−_) (Figure 4A).

**Figure 4.**
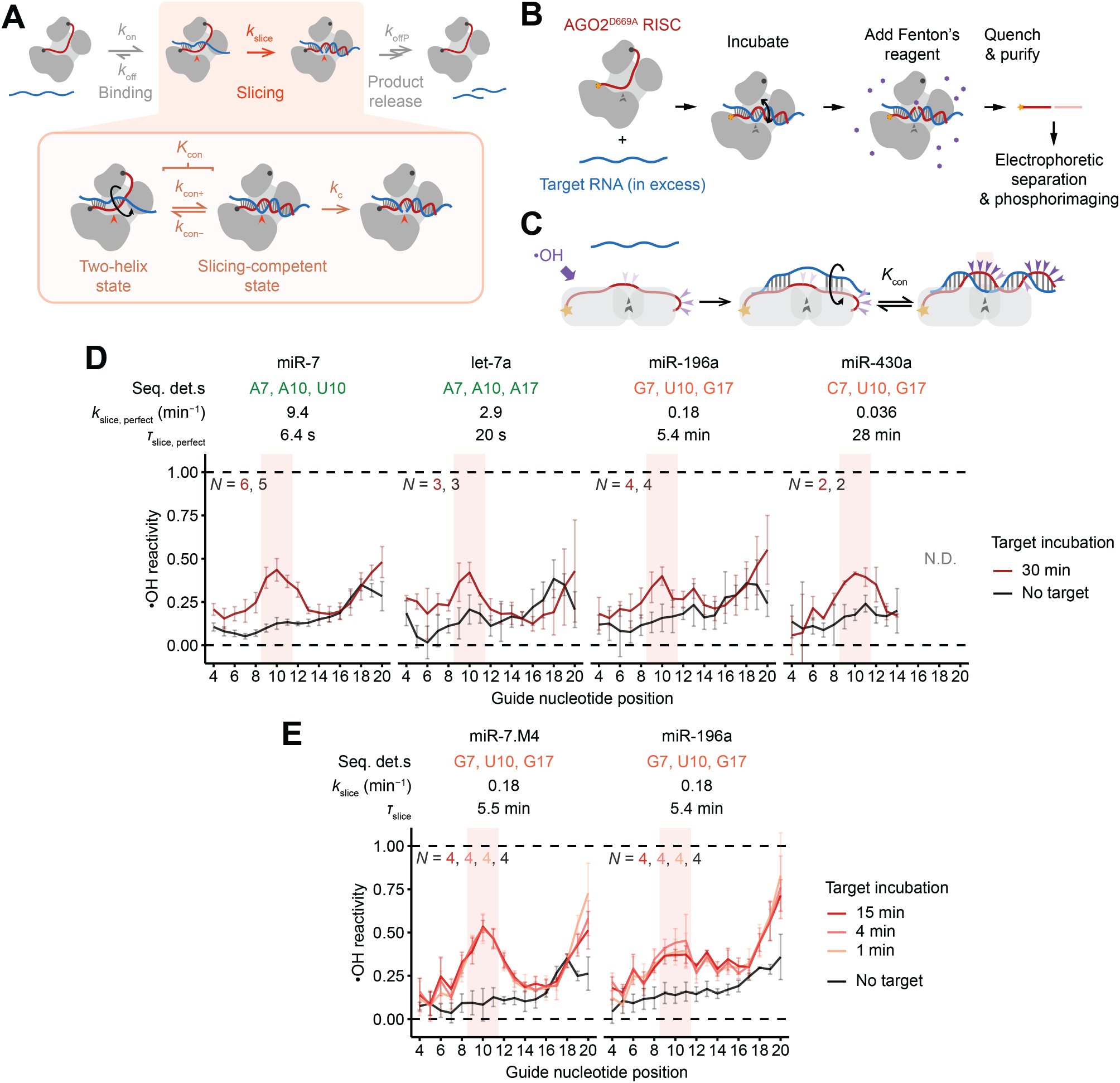
Formation of central pairing does not limit *k*_slice_ of fully complementary targets. (A) Expanded kinetic scheme illustrating how conformational dynamics of the RISC−target complex could influence slicing kinetics; otherwise, as in Figure 1B. Under thermodynamic conformational control, *k*_slice_ is a function of *K*_con_ (i.e., *k*_con+_/*k*_con–_); under kinetic conformational control *k*_slice_ is a function of *k*_con+_. If not under conformational control, *k*_slice_ is under control of *k*_c_. (B) Schematic of the hydroxyl-radical footprinting experiment. ^32^P label of the guide is indicated with a yellow star, and the D669A mutation is represented as a broken gray caret; otherwise colors are as in (A). (C) Schematic of changes in solvent accessibility of the guide upon conformational change to the slicing-competent state, as probed by hydroxyl radicals. Exposed guide backbone positions predicted to be readily cleaved by hydroxyl radicals are indicated with purple carets. Otherwise, colors are as in (B). (D) Evidence that thermodynamics of conformational change do not influence slicing rates of fully complementary targets. Footprinting reactivity values measured after RISC was bound to either no target or fully complementary target are plotted for each of four guide RNAs spanning the range of *k*_slice_ for perfectly complementary targets (Figure 2D). Reactivity values were normalized to those of naked-guide or quenched-reagent samples (dashed lines). The band for cleavage after position 21 was not cleanly separated from the full-length band, and bands for cleavage after positions 1−3 were not resolved from the salt front. Cleavage after positions 15−20 of miR-430a could not be confidently quantified (N.D.) due to minor trimmed isoforms of the guide in the input (Figure S6B). Error bars indicate 95% CIs. Positions 9−11 are shaded. Number of replicates for each condition is shown as *N* in the corresponding color. (E) Evidence that kinetics of conformational change do not influence slicing rates of fully complementary targets. Footprinting reactivity values measured after RISC was incubated with fully complementary target for the indicated amount of time are plotted for each of two guide RNAs with relatively slow *k*_slice_ values; otherwise, as in (D). See also Figure S6.

With this model in mind, we used hydroxyl-radical footprinting to measure the extent to which different RISC–target complexes populate the centrally paired configuration. In this chemical-footprinting procedure, hydroxyl radicals cleave the RNA backbone at solvent-exposed positions, creating a gel banding pattern that reports on solvent accessibility of each backbone position of an end-labeled RNA (Figures 4B and 4C).^64^ For this analysis, RISCs were generated using AGO2^D669A^ (containing an active-site substitution designed to prevent slicing^9,41^) and each of four end-labeled miRNAs, chosen to span the breadth of *k*_slice_ values.

For each complex, footprinting was performed after incubation with either no target or one of several target RNAs, each with different pairing potential to the miRNA (Figures 4B and S6A). After incubation with no target, miRNA backbones had low reactivity across their lengths, with a moderate uptick near position 18 (Figures 4D and S6B), as expected from solvent accessibilities determined from crystal structures^33–35^ (Figure 4C). After incubation with targets that paired to only the seed or to the seed with five additional supplementary pairs, the reactivity profiles were largely unchanged (Figures S6B and S6C), in agreement with crystal structures.^36,41^ After incubation with targets with seed and extensive 3′ base-pairing, backbones of some miRNAs acquired increased solvent accessibility from around position 12 to the 3′ end (Figures S6B−D), again in agreement with structures.^42^ After incubation with perfectly complementary target RNAs, the reactivity profiles acquired strong peaks at positions 9−11 and after position 18, which closely matched those expected for the slicing-competent state (Figures 4C, 4D, and S6B). Importantly, the profiles observed for each of the four miRNAs were similar, despite their >250-fold range in *k*_slice_ values, in stark contrast to the expectation that steady-state occupancy of the centrally paired, slicing-competent state is lower for guides with slower *k*_slice_. Indeed, across the four miRNAs, we found no significant differences in the heights of the central-region reactivity peaks diagnostic of the slicing-competent state, implying no detectable differences in the equilibrium occupancies of the slicing-competent conformation (*K*_con_) for fast- and slow-slicing guides.

Having acquired evidence disfavoring the thermodynamic model, we investigated the possibility of a kinetic model, in which slow initial formation of the slicing configuration (i.e., slow *k*_con+_) limits *k*_slice_. In this model, slow-slicing guides populate the centrally paired, slicing-competent configuration more slowly, at the same rate at which their targets are sliced. To test this model, we probed two slow-slicing guides (miR-196a and miR-7.M4) after incubating each with its fully complementary target for times (1, 4, or 15 min) that bracketed their *τ*_slice_ values (5.4 and 5.5 min, respectively). For both guides, no discernible difference was observed across the different times (Figures 4E and S6B).

We conclude that the large conformational change required to form the centrally paired RISC−target ternary complex does not limit the kinetics of slicing perfectly complementary targets, either kinetically or thermodynamically. The centrally paired conformation forms within a few minutes, even for slow RISCs, and surprisingly, at equilibrium, it is no more populated for fast RISCs than slow RISCs. Presumably the sequence determinants operate downstream of this large conformational change, producing subtle structural differences in the slicing-competent state that alter rate-limiting *k*_c_, and consequently *k*_slice_, but are too small to detect by chemical footprinting.

### Weak pairing at positions 6−7 appears to facilitate a favorable helical distortion

We next sought to identify the structural basis by which sequence determinants might influence slicing kinetics, with particular interest in the determinants at positions 7 and 17, which were distant from the cleavage site. With respect to position 7, we hypothesized that enhanced slicing associated with an A:U or U:A pair might be imparted through the same mechanism as the enhanced slicing previously observed for a G:G mismatch at position 6,^48^ and that this unknown mechanism might involve weak pairing at positions 6 or 7. To test this hypothesis, we investigated how enhancement by a position-6 G:G mismatch was affected when the pair at position 7 was changed. In the context of miR-7.M3, which makes a disfavored G:C base-pair at position 7, introducing a G:G mismatch at position 6 increased *k*_slice_ by 2.4-fold (Figures 5A and 5B). In contrast, this enhancement was absent in the context of miR-7, which already makes a favored weak A:U base-pair at position 7. Moreover, when comparing slicing directed by miR-7 and miR-7.M3, the effect from the position-7 substitution diminished from 4.4-fold to 1.9-fold in the context of the G:G position-6 mismatch (Figures 5A and 5B). Thus, enhancement of *k*_slice_ by the weak pair at position 7 and by the G:G mismatch at position 6 were non-additive and at least partially redundant, as expected if they were mediated through the same mechanism. In contrast, an analogous comparison using substitutions at positions 10−11 found no significant impact on the enhancement of *k*_slice_ by the position-6 mismatch, as expected if the determinants were acting independently (Figures S7A and S7B).

**Figure 5.**
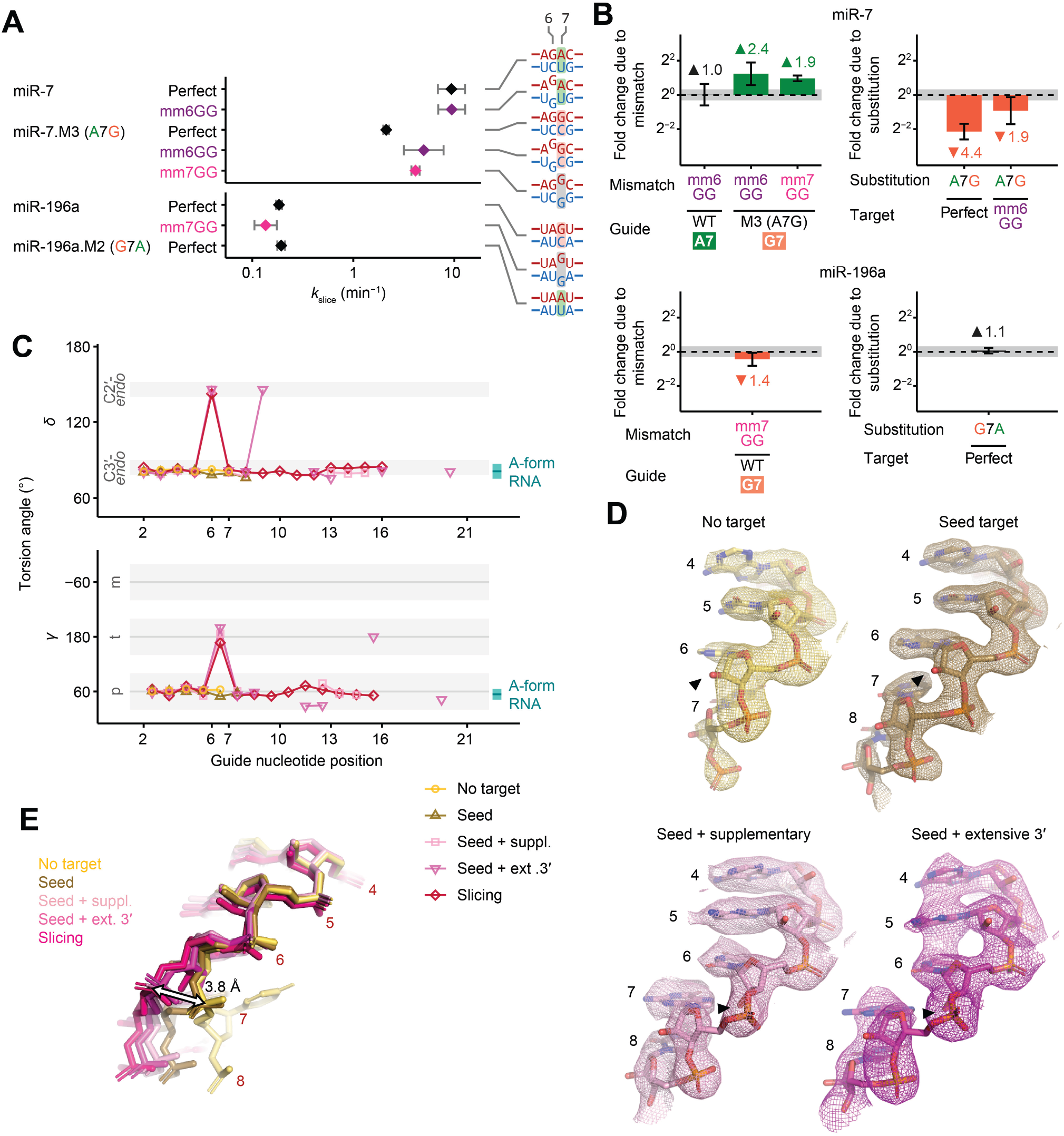
Enhancement of slicing by weak pairs or mismatches at positions 6−7 aligns with local backbone distortion. (A) Effects of mismatches at positions 6–7. Schematics on the right illustrate pairing at positions 5−8; (B) otherwise, as in Figure 2C. Values with perfectly complementary targets are replotted from Figure 2C. (B) Comparisons of *k*_slice_ values observed in (A), plotted in log-scale. Left panels show the fold change in *k*_slice_ due to different mismatches in the target, in the context of different guide RNAs. Nucleotide identity at the position-7 sequence determinant is indicated for each guide. Right panels show the fold change in *k*_slice_ due to substitutions introduced at position 7, in the context of either perfectly complementary targets or targets harboring a mismatch at position 6. Fold-change values are also indicated above or below the bar plots. Error bars indicate 95% CIs after propagating uncertainty from model-fitting for the two *k*_slice_ values compared. Colors are as in (A) and Figure 2C. 95% CI of expected background variation is indicated with gray shading (Figure S5A). (C) Changes in *δ* and *γ* torsion angles observed in the guide backbone as RISC assumes different conformational states. Measurements are from structure models in which the guide RNA has either no target (4W5N, yellow^36^), seed-pairing (4W5R, brown^36^), seed and supplementary-pairing (6N4O, light pink^41^), seed and extensive 3′-pairing (6MDZ, magenta^42^), or full pairing from position 2 to 16 (7SWF, deep pink^43^). Values are only shown for base-paired nucleotides or, in the case of the no-target state, positions 2−7, which are pre-organized into an A-form-like helix by AGO. Six values corresponding to unexpected conformation outliers, indicating possible poor model-fit, were found in positions 13−19 of states with either seed and supplementary pairing or seed and extensive 3′-pairing; these were not plotted (Figure S7D). Expected ranges of angles for canonical conformers are shown in gray, and expected ranges of angles for A-form helices are indicated in turquoise on the right.^80^ (E) Distortion of the guide backbone as RISC assumes different conformational states. Shown are structure models of the guide backbone at positions 4−8 for each of the structures in (C) solved using crystallography, overlaid with electron density omit maps (2*mF*_o_ − *DF*_c_)^81^ contoured at 1.6 *σ*, shown in mesh. The 2′-hydroxyl of position 6 is indicated with a black triangle for each model. Position 8 in the no-target state was not resolved and thus is not shown. Colors are as in (C). (E) As in (D) but for all five structure models and without the electron density maps. Models were overlaid and aligned at AGO MID and PIWI domains. Displacement of the position 7 phosphate observed as RISC shifts from the no-target state to the slicing state is indicated with an arrow. See also Figure S7 and Table S1.

To further test the idea that the weak pair at position 7 and the G:G mismatch at position 6 were not exerting their influence through separate mechanisms, we tested target RNA substrates with a G:G mismatch at position 7. A G:G mismatch at position 7 of miR-7.M3 enhanced *k*_slice_ by 1.9-fold, similar to its effect at position 6 (Figures 5A and 5B). In comparison, a G:G mismatch at position 7 of miR-196a did not enhance *k*_slice_, which resembled the lack of effect from mutating a G:C base-pair to an A:U base-pair at position 7 of miR-196a (Figures 5A, 5B, and 2C), providing further evidence that the enhancement of *k*_slice_ by the position-7 sequence determinant was related to the enhancement by G:G mismatches at positions 6 and 7, as expected if they were mediated through the same mechanism.

We hypothesized that slicing might require a structural distortion between positions 6 and 7, which a weaker or disrupted base-pair might favor. To investigate this idea, we analyzed five high-resolution structure models of slicing-competent AGO proteins, including four of human AGO2 and one of *Arabidopsis thaliana* AGO10. These included structures with either 1) no target,^36^ 2) a target paired to only the seed region,^36^ 3) a target paired additionally to positions 13−16,^41^ 4) a target paired more extensively to the 3′ half of the guide,^42^ or 5) a target paired perfectly to positions 2–16.^43^ Examining the torsion angles of the guide-RNA backbone in these structure models revealed a striking difference in the three structures with pairing beyond the seed; these three each had a clear local deviation from an A-form helix between positions 6 and 7 (Figures 5C, S7C, and S7D). Although most well-resolved nucleotides in the miRNA had geometries corresponding to a C3′-*endo* sugar pucker, typical of A-form RNA, for these miRNAs paired beyond the seed, unusual backbone geometry at position 6 implied a distinct C2′-*endo* sugar pucker, which was evident from deviated *δ* and *ν*_*2*_ torsion angles and shifted 2′-hydroxyl groups in electron density maps (Figures 5C, 5D, and S7C). Consequently, the RNA backbone adopted a kink between positions 6 and 7, as reflected in the distinct γ torsion angles and conformer classes (Figures 5C and S7D), and this distortion resulted in a 3.8-Å shift in the position of the guide at position 7 (Figure 5E). Based on these observations, we posit that this distortion is necessary for the correct positioning and register of the guide−target helix for base-pairing and slicing, and that it may be favored by a weak or mismatched base-pair at positions 6 or 7. Supporting this model, incorporating a flexible (*S*)-glycol nucleic acid (*S*-GNA) backbone modification at position 6 or 7 but not at any other positions of the guide or passenger strand enhances efficacy of siRNAs in cells.^65,66^ Indeed, authors of the *S*-GNA studies note that the beneficial effect of *S*-GNA corresponds to the site of distortion observed in the structures. Together, our studies suggest that favoring this distortion acts to increase *k*_slice_, which presumably occurs for the optimized *S*-GNA-containing siRNA currently in clinical trials.^67^ A similar backbone distortion was also observed in high-resolution models of prokaryotic AGO proteins (Figures S7E and S7F),^68^ suggesting that it is an ancestral feature shared by diverse AGO homologs.

### Rapid slicing of terminally mismatched sites appears to require strong central pairing

Preference for a weak pair at position 17 prompted examination of slicing kinetics of targets harboring mismatches at this position. In contrast to mismatches at positions 6 or 7, mismatches at position 17 strongly disrupted slicing, leading to an 8.3-to 64-fold reduction of *k*_slice_ in the six cases examined (Figure 6A). Thus, in contrast to the case of position 7, preference for an A:U/U:A pair at position 17 did not seem to stem from a preference for weak pairing.

**Figure 6.**
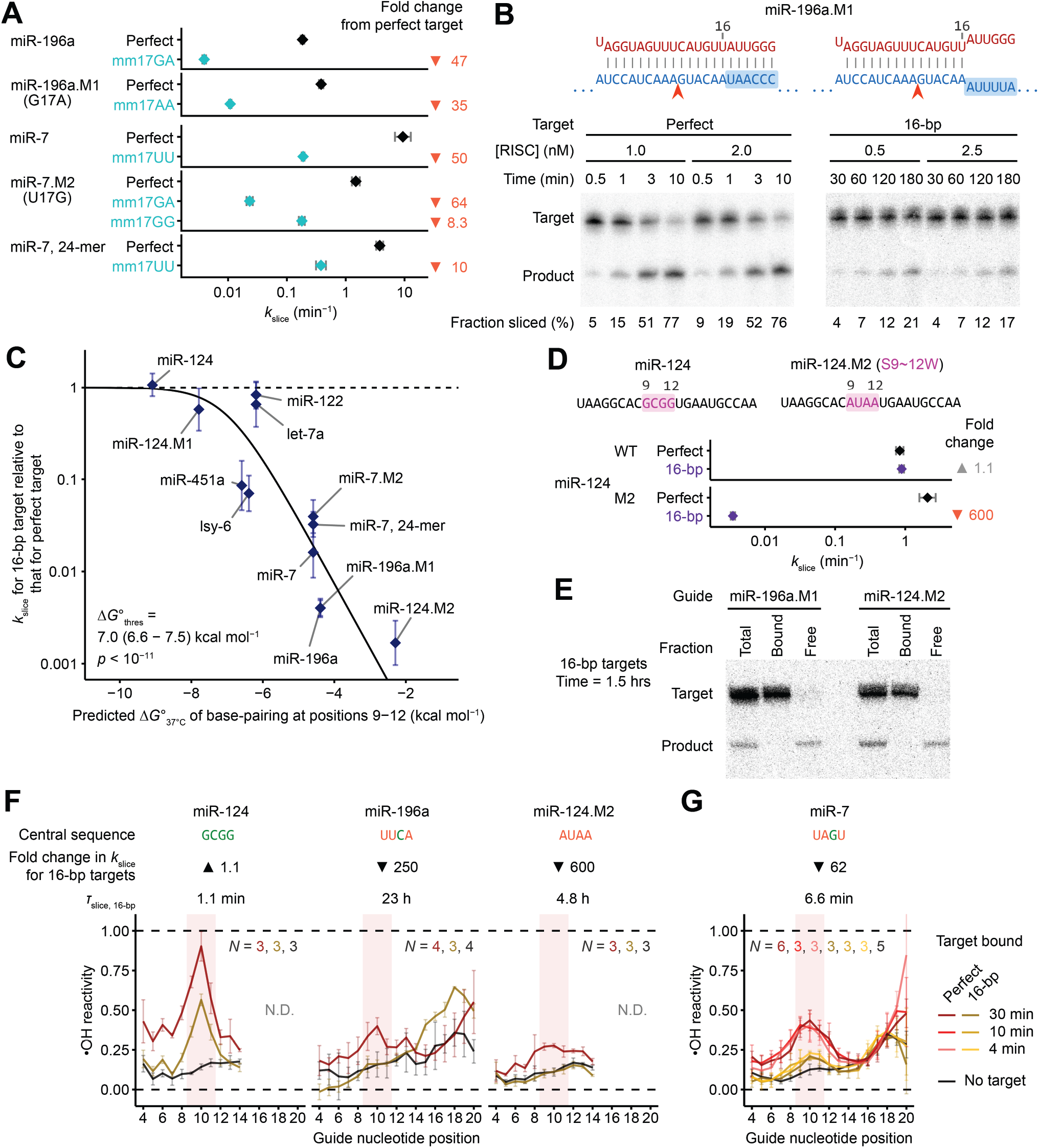
Strong central pairing compensates for terminal mismatches, which otherwise reduce the fraction of complexes in the slicing configuration. (A) The importance of pairing to position 17. Measured are *k*_slice_ values for perfectly complementary targets (black) or targets harboring a mismatch at position 17 (cyan); otherwise, as in Figure 2C. Values with perfectly complementary targets are replotted from Figure 2C. (B) The importance of pairing to positions 17–22. Shown are pairing diagrams and representative results for miR-196a.M1-directed slicing of its perfectly complementary and 16-bp targets. Target nucleotides changed to introduce terminal mismatches are highlighted in blue; otherwise, as in Figure 2B. (C) Relationship between fold change in *k*_slice_ observed for 16-bp targets and predicted base-pairing energy at positions 9−12. Error bars indicate 95% CIs based on propagated uncertainty from model-fitting for the two *k*_slice_ values considered. Curve shows the nonlinear least-squares best fit of a thermodynamic equation involving one fitted parameter (Δ*G*°_thres_); its fitted value is shown with its 95% CI and *p* value as calculated by a *t*-test. (D) Mutational evidence supporting the relationship between predicted strength of central pairing and the effect of terminal mismatches. Measured are *k*_slice_ values for perfectly complementary or 16-bp targets, for miR-124 and its mutant (miR-124.M2), in which the central G/C nucleotides of miR-124 were replaced with A/U nucleotides. Substituted positions are highlighted in the schematic. Values from 16-bp targets are plotted in purple; otherwise, as in (A). Values with perfectly complementary targets are replotted from Figure 2D. (E) Evidence that slowly sliced 16-bp targets are nonetheless fully bound. Shown are results of slicing assays with 16-bp targets, in which the RISC-bound and free RNA species were separated by filter binding after 1.5 hours of incubation. RISC and target were initially at 1.0 and 0.05 nM, respectively. (F) Evidence that forming the centrally paired conformation limits *k*_slice_ when guides with weak central pairing are slicing targets with terminal mismatches. Footprinting reactivity values observed when RISC was bound to no target, 16-bp target, or perfectly complementary target, are plotted for each of three miRNAs. Results for miR-196a with no target and with perfectly complementary target are replotted from Figure 4D. Positions 15−20 of miR-124 and miR-124.M2 could not be confidently quantified (N.D.) due to minor trimmed isoforms of the guide in the input (Figure S6B). Otherwise, as in Figure 4D. (G) Evidence that the conformational change thermodynamically limits *k*_slice_ when guides with weak central pairing are slicing targets with terminal mismatches. This panel is as in Figure 4E, except it shows reactivity values for miR-7 RISC as it engages with either perfect or terminally mismatched target. Results for miR-7 RISC after incubation with either no target or for 30 min with perfectly complementary target are replotted from Figure 4D. See also Figure S6 and Table S1.

The requirement for pairing at position 17 seemed at odds with reports that pairing beyond position 16 is dispensable for efficient slicing.^45,47^ Alternatively, because previous studies had examined pairing requirements for only two miRNAs (let-7a and miR-21), which differed from the ones we examined, pairing beyond position 16 might be dispensable for efficient slicing guided by some sequences but not others. To investigate this possibility, we measured *k*_slice_ values of targets with complementarity up to only position 16 (referred to as 16-bp targets) for 12 different guide RNAs, and compared them to *k*_slice_ values of perfectly complementary targets. Although *k*_slice_ was reduced only minimally for some guide RNAs, it was reduced by >100-fold for others (Figures 6B and 6C). This variation was strongly correlated with predicted pairing energy of positions 9–12 of the guide−target RNA duplex (Figure 6C), which is proposed to be the final region of the guide−target duplex to form prior to slicing.^2^ For example, the miR-124 complex, which had very stable predicted central pairing, imparted by GCGG at positions 9−12, had a *k*_slice_ value for its 16-bp target that was indistinguishable from that observed for its perfectly complementary target. In contrast, the miR-196a.M1 complex, which had UUCA at positions 9−12, sliced its 16-bp target 250-fold slower than it sliced its perfectly complementary target (Figure 6B). Changing positions 9−12 of miR-124 from GCGG to AUAA (miR-124.M2) imparted a 600-fold drop in *k*_slice_ when slicing the 16-bp target, which took 4.8 h (95% CI, 4.5–5.1 h), but imparted little effect when slicing a perfectly complementary target (Figures 2D, 6C, and 6D).

To confirm that this drop represented a change in *k*_slice_ and not target association, we used filter binding to monitor association of slow-slicing targets at pre-steady-state timepoints. For both miR-196a.M1 and miR-124.M2, essentially all of the unsliced 16-bp target was bound by RISC at these early time points, confirming that the slow reaction rates were due to slow slicing (Figure 6E).

We conclude that base-pairing beyond position 16 is dispensable for efficient slicing only when the central region has high predicted pairing stability. This pairing energy is likely required to drive the conformational equilibrium to the centrally paired configuration at the expense of disrupting favorable contacts between the guide-RNA 3′ region and AGO2, including contacts with the PAZ domain—contacts that for fully complementary sites would otherwise be disrupted by target pairing.^42^ Thus, GC content at positions 9−12 constitutes an additional *k*_slice_ determinant, but one that applies specifically to 3′-mismatched targets.

### Formation of continuous helix thermodynamically limits slicing of terminally mismatched targets

Our proposed model for the ability of strong central pairing to favor the slicing configuration of terminally mismatched targets predicted that guide RNAs with weak central pairing would populate the slicing configuration more poorly when associated with 3′-terminally mismatched (16-bp) targets than when associated with fully complementary targets. Hydroxyl radical footprinting confirmed this prediction. As expected from the earlier footprinting (Figure 4D), all three complexes tested had elevated reactivity at positions 9–11 (diagnostic of achieving the slicing-competent conformation) after engaging fully complementary targets (Figures 6F and S6B).

However, only the complex with strong predicted central pairing (miR-124) had elevated reactivity at positions 9−11 after engaging its 16-bp target; guides with weak predicted central pairing (miR-196a and miR-124.M2) had reactivity profiles closely resembling the baseline of initial-binding states (Figures 6F and S6B).

To determine if this difference was thermodynamic or kinetic, we probed miR-7 engaged with its 16-bp target, which had an intermediate rate of slicing amenable to an incubation time course of 4–30 min. Across this time course, no discernible difference in the reactivity profiles was found, indicating that for guides with weak central pairing, populating the slicing-competent conformation with a 3′-mismatched target is thermodynamically, not kinetically, limited (Figures 6G and S6B).

## DISCUSSION

Our study reveals the interplay between guide-RNA sequence, RISC conformation, and slicing activity (Figure 7A). The discovery of significant variation in slicing kinetics when AGO2 is loaded with different guides emphasizes that different miRNAs/siRNA sequences can have substantially different biochemical behaviors. Some of the biochemical properties of AGO2, such as slicing between nucleotides paired to positions 10−11,^3^ appear generalizable to all guide RNAs. Other properties, however, such as relative strengths of different canonical and noncanonical site types,^49^ positional preferences of additional 3′ base-pairing,^69^ kinetics of slicing (Figure 2D), tolerance of mismatches for slicing (Figure 6C), and the rate-limiting steps of multiple-turnover slicing (Figures 1D, S5D, and S5E), can substantially vary, depending on the guide.

**Figure 7.**
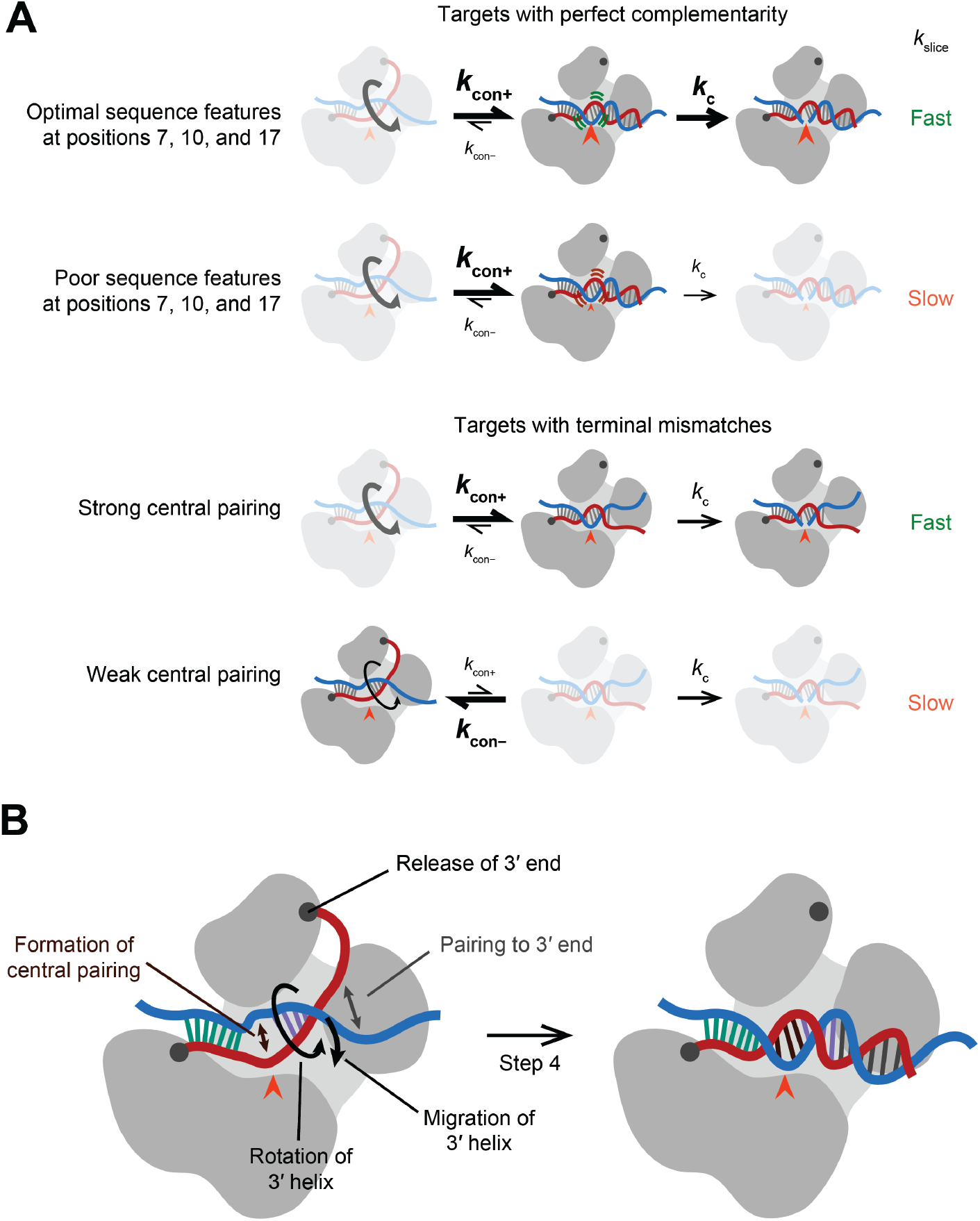
Sequence and conformational determinants of slicing by AGO2. (A) Schematic of the proposed relationships between guide-RNA sequence determinants, conformational dynamics, and *k*_slice_ for either perfectly or partially complementary targets. When RISC is bound to perfectly complementary targets, the slicing-competent conformation is always well-populated. RNA sequence determinants appear to confer subtle structural changes that affect the efficiency of chemistry (*k*_c_). When RISC is bound to targets mismatched after position 16, only guides with strong central pairing fully populate the centrally paired, slicing-competent configuration. (B) The concerted, mutually reinforcing movements proposed to occur as target-bound RISC reaches the slicing conformation.

Guide-specific properties were illustrated most emphatically by the 600-fold range in 3′-mismatch tolerance. The striking relationship between the strength of central pairing and 3′-mismatch tolerance supports the notion that the multiple movements occurring during the final step of the four-step process of pairing to a fully complementary target are interconnected. Indeed, we propose that all of the movements at this final step, in which the two-helix conformation transitions to the fully paired conformation, enable or promote each other (Figure 7B). In this view, propagation of pairing from the second helix towards the guide 3′ terminus promotes (and is promoted by) release of the guide 3′ terminus from the PAZ domain to untether the second helix, which enables (and is promoted by) rotation and repositioning of the second helix, which enables (and is promoted by) pairing in the central region. Such interconnected movements provide the simplest explanation for how the strength of pairing to the central region of the guide can modulate 3′ mismatch tolerance.

The influence of terminal pairing on the equilibrium between the two-helix and the centrally paired conformations helps explain why an attempt to crystallize the centrally paired, slicing-competent conformation of miR-122 RISC was unsuccessful.^41^ This attempt used a 16 nt target. When bound to such a short target, the complex would occupy the two-helix conformation more frequently than when bound to a longer, fully complementary target. Thus, some occupancy of the two-helix conformation (even if it does not predominate for miR-122; Figure 6C), together with the influence of crystal contacts, could lead to inadvertent crystallization of the two-helix conformational intermediate.

Our insights into target slicing presumably also apply to many cases of RISC maturation. Although most miRNA–passenger strand duplexes contain mismatches that enable AGO to eject the passenger strand without slicing it, at least two human miRNAs require slicing for their maturation within AGO2.^18–21^ Most synthetic and endogenous siRNA duplexes are also highly complementary and presumably require slicing for the passenger-strand removal.^22–24^

The degree to which our results extend beyond eukaryotic AGO proteins is unclear. For example, prokaryotic AGO from *Thermus thermophilus* and eukaryotic PIWI each have a wider central cleft, with evidence that pairing can propagate directly from the seed, without nucleation of a second helix.^32,70,71^ Nonetheless, these proteins might still prefer the two-helix pathway for some guide RNAs, such as those with weak or imperfect central pairing.

When examining different guide-RNA sequences, the dynamic ranges of the forward kinetic parameters (*k*_on_, *k*_slice_, and *k*_offP_) overlap each other extensively (Figure S5E), such that any of these steps can become rate-limiting, and therefore potentially influence RNAi efficacy. For example, a guide with *τ*_slice_ of 12 min would not be able to achieve more than 94% knockdown of an mRNA with a half-life of 2 h (the median half-life for mRNAs in 3T3 cells^72^) and no more than 78% knockdown of an mRNA with a half-life of 30 min.

The range of *k*_slice_ values also spans typical kinetics of other cellular functions, such as translation, although increasing the number of ribosomes translocating through the target site did not appear to influence reporter repression. One explanation for our inability to detect an effect of ribosomes centers on the possibility that strong affinity of RISC for perfectly complementary sites^39^ might cause ribosomes to stall before displacing RISC^73^ and thereby increase the incidence of ribosome collisions that trigger the no-go decay pathway, resulting in the degradation of the mRNA,^74^ as observed for antisense oligonucleotides paired to coding sequences.^75^ Thus, inducing translation of the reporter might have had two offsetting effects: increased ribosome-mediated displacement of RISC, causing reduced RISC-catalyzed slicing, and increased ribosome collisions, causing increased degradation through no-go decay. Other potentially confounding pathways include TNRC6-mediated deadenylation and decapping of slowly sliced target transcripts^1^ and target-directed degradation of the bound RISC.^76,77^

We propose that the sequence determinant at position 7 favors a helical distortion associated with more extensively paired conformations, but potential mechanistic underpinnings of determinants at positions 10 and 17 are less clear. The determinant at position 10 was the strongest of the three and coincided with the strongest sequence-specific feature found in siRNA functional screens.^56^ Its proximity to the slicing site suggests that it might form favorable interactions with the central chamber of AGO2 in the slicing-competent conformation, allowing the optimal configuration of the active site to be more readily adopted. With respect to the determinant at position 17, additional insight awaits high-resolution structural studies of the fully paired slicing conformation.

The observation that flexible or mismatched pairs at position 6–7 of the guide-RNA seed promote *k*_slice_ illustrates that some imperfections in seed pairing can be favored in this context in which extensive pairing to the remainder of the guide RNA are more than sufficient for target binding. Of course, in the context of typical miRNA–target interactions, which are primarily mediated by seed pairing, these imperfections in the seed match greatly reduce binding and repression.^49,65^

Our results are relevant for designing siRNAs for research and therapeutic applications. First, our results support design recommendations proposed two decades ago regarding positions 7, 10, and 17.^56^ Moreover, our results provide a rationale for these guidelines, indicating that nucleotide identity at these positions affects *k*_slice_, as opposed to affecting siRNA intracellular uptake, loading, stability, target binding, or product release. Second, our results speak to the importance of pairing beyond siRNA position 16. Some have suggested that siRNAs with complementarity ending at position 16 would have enhanced performance.^47^ Our results indicate that this would be a viable strategy only for siRNAs with strong pairing at their central region. Moreover, our results show that designs involving weaker central pairing would increase specificity for sites that fully pair to the miRNA 3′ region, thereby reducing off-target slicing at sites with only partial complementarity. The striking differences observed for tolerance of 3′-mismatches also indicate that initial structural studies of the slicing configuration, which use targets that do not pair beyond position 16,^41,43^ are most relevant for guides with very strong central pairing, whereas structural studies with fully complementary targets will be most relevant for studying the slicing configuration of guides with weaker central pairing.

### Limitations of the study

When including all the mutant guide RNAs, we purified and kinetically examined RISCs loaded with 29 different guide RNAs (Table S1)—a large increase over the number of complexes previously examined, but insufficient to exhaustively identify *k*_slice_ sequence determinants by regression. Further assessment of more guide sequences will presumably reveal additional determinants and enable interdependencies between determinants to be identified, ultimately generating an algorithm for accurately predicting slicing rates based on guide sequences.

## ACKNOWLEDGEMENTS

We thank S. E. McGeary, D. H. Lin, T. M. Pham, K. Xiang, and others in the Bartel laboratory for helpful discussions. We thank S.E.M., T.M.P., D. Briskin, and M. H. Hall for several purified RISCs. We thank the Whitehead Proteomics Core for mass spectrometry analyses. This work was supported by NIH grant GM118135. D.P.B. is an investigator of the Howard Hughes Medical Institute.

## AUTHOR CONTRIBUTIONS

P.Y.W. and D.P.B. conceived the project, designed the study, and wrote the manuscript. P.Y.W. performed the experiments and data analyses.

## DECLARATION OF INTERESTS

D.P.B. has equity in Alnylam Pharmaceuticals, where he is a co-founder and advisor. D.P.B. is a member of Molecular Cell’s advisory board. P.Y.W. declares no competing interests.

## STAR METHODS

### KEY RESOURCES TABLE

| REAGENT or RESOURCE | SOURCE | IDENTIFIER |
| --- | --- | --- |
| Bacterial and virus strains |  |  |
| One Shot TOP10 Chemically Competent *E. coli* | Thermo Fisher | Cat#C404003 |
| MAX Efficiency DH5α Competent Cells | Thermo Fisher | Cat#18258012 |
| One Shot OmniMAX 2 T1R Chemically Competent *E. coli* | Thermo Fisher | Cat#C854003 |
| Chemicals, peptides, and recombinant proteins |  |  |
| SUPERase•In | Invitrogen | Cat#AM2694 |
| Thermostable Inorganic Pyrophosphatase | New England Biolabs | Cat#M0296 |
| T7 RNA Polymerase | Purified in-house | N/A |
| RQ1 DNase | Promega | Cat#M6101 |
| Gel Loading Buffer II | Invitrogen | Cat#AM8547 |
| Shrimp alkaline phosphatase | New England Biolabs | Cat#M0371 |
| Quick CIP | New England Biolabs | Cat#M0525 |
| T4 Polynucleotide Kinase | New England Biolabs | Cat#M0201 |
| *Vaccinia* capping system | New England Biolabs | Cat#M2080 |
| [γ-<sup>32</sup>P] ATP | PerkinElmer | Cat#NEG035C001MC |
| [α-<sup>32</sup>P] GTP | PerkinElmer | Cat#NEG006H250UC |
| Lipofectamine 2000 | Invitrogen | Cat#11668019 |
| cOmplete Mini EDTA-free Protease Inhibitor Cocktail | Roche | Cat#11836170001 |
| Dynabeads MyOne Streptavidin C1 | Invitrogen | Cat#65002 |
| Anti-FLAG M2 magnetic beads | Millipore | Cat#M8823 |
| IGEPAL CA-630 | Sigma-Aldrich | Cat#I3021 |
| BSA | New England Biolabs | Cat#B9000 |
| Yeast tRNA | Life Technologies | Cat#AM7119 |
| 3xFLAG peptide | Sigma-Aldrich | Cat#F4799 |
| Proteinase K | Invitrogen | Cat#25530049 |
| Phenol:chloroform:isoamyl alcohol (25:24:1) | Sigma-Aldrich | Cat#P2069 |
| RNase T1 | Invitrogen | Cat#AM2283 |
| Ammonium iron(II) sulfate | Sigma-Aldrich | Cat#09719 |
| Hydrogen peroxide, 30% | Sigma-Aldrich | Cat#216763 |
| Sodium sulfite | Sigma-Aldrich | Cat#S0505 |
| Desferroxamine mesylate | Sigma Aldrich | Cat#D9533 |
| Ferric ammonium citrate | Sigma Aldrich | Cat#F5879 |
| Critical commercial assays |  |  |
| Micro Bio-Spin P-30 columns | Bio-Rad | Cat#7326250 |
| Micro Bio-Spin P-6 columns | Bio-Rad | Cat#7326221 |
| KAPA HiFi HotStart Ready Mix | Roche | Cat#07958935001 |
| NEBuilder HiFi DNA Assembly Master Mix | New England Biolabs | Cat#E2621 |
| Dual-Luciferase Reporter assay system | Promega | Cat#E1960 |
| PureLink HiPure Expi Plasmid Megaprep Kit | Invitrogen | Cat#K210008XP |
| Deposited data |  |  |
| Raw and analyzed data | This paper | Mendeley Data, DOI: 10.17632/j82nkmfppj.1 |
| Crystal structure model of AGO2 (no target) | Schirle et al.<sup>36</sup> | PDB: 4W5N |
| Crystal structure model of AGO2 (seed-match only) | Schirle et al.<sup>36</sup> | PDB: 4W5R |
| Crystal structure model of AGO2 (seed + supplementary pairing) | Sheu-Gruttadauria et al.<sup>41</sup> | PDB: 6N4O |
| Crystal structure model of AGO2 (seed + extensive 3' pairing) | Sheu-Gruttadauria et al.<sup>42</sup> | PDB: 6MDZ |
| Cryo-EM structure model of AtAGO10 (slicing-competent state) | Xiao et al.<sup>43</sup> | PDB: 7SWF |
| Crystal structure model of TtAgo (slicing-competent state) | Sheng et al.<sup>68</sup> | PDB: 4NCB |
| Experimental models: Cell lines |  |  |
| Human: HEK293FT cells | Bartel laboratory | N/A |
| Human: HEK293T cells | Bartel laboratory | N/A |
| Human: HeLa cells | Bartel laboratory | N/A |
| Oligonucleotides |  |  |
| See Table S2 | This paper | N/A |
| Recombinant DNA |  |  |
| pMaxGFP | Lonza | N/A |
| pcDNA3.3-3xFLAG-AGO2 | Addgene | Addgene ID: 136687 |
| pcDNA3.3-3xFLAG-AGO2(D669A) | This study | Addgene ID: 220461 |
| pcDNA3.3-3xFLAG-AGO2(S387A) | This study | Addgene ID: 220462 |
| pcDNA3.3-3xFLAG-AGO2(S387D) | This study | Addgene ID: 220463 |
| pSliceRep_A_miR-7 | This study | Addgene ID: 220464 |
| pSliceRep_A_miR-7.M4 | This study | Addgene ID: 220465 |
| pSliceRep_A_miR-196a | This study | Addgene ID: 220466 |
| pSliceRep_A_miR-196a.M3 | This study | Addgene ID: 220467 |
| pSliceRep_B_miR-7 | This study | Addgene ID: 220468 |
| pSliceRep_B_miR-7.M4 | This study | Addgene ID: 220469 |
| pSliceRep_B_miR-196a | This study | Addgene ID: 220470 |
| pSliceRep_B_miR-196a.M3 | This study | Addgene ID: 220471 |
| Software and algorithms |  |  |
| R (v4.3.3) | The R Foundation | https://cran.r-project.org/ |
| Python (v3.11.4) | Python Software Foundation | https://www.python.org/ |
| Levenberg-Marquardt algorithm, from minpack.lm package | Timur V. Elzhov, Katharine M. Mullen, Andrej-Nikolai Spiess, Ben Bolker | https://cran.r-project.org/package=minpack.lm |
| Livermore solver for ordinary differential equations (LSODE), from deSolve package | Soetaert et al.<sup>82</sup> | https://cran.r-project.org/package=deSolve |
| modFit data-fitting algorithm, from FME package | Soetaert et al.<sup>83</sup> | https://cran.r-project.org/package=FME |
| Multi Gauge | Fujifilm | N/A |
| ViennaRNA 2.0 package | Lorenz et al.<sup>84</sup> | https://www.tbi.univie.ac.at/RNA/ |
| ImageQuant TL | Cytiva | v10.2 |
| ImageJ 1.53n | National Institutes of Health | https://imagej.net/ij/ |
| Proteome Discoverer | Thermo Fisher | https://www.thermofisher.com/us/en/home/industrial/mass-spectrometry/liquid-chromatography-mass-spectrometry-lc-ms/lc-ms-software/multi-omics-data-analysis/proteome-discoverer-software.html |
| Mascot | Matrix Science | https://www.matrixscience.com/ |
| Scaffold 5 | Proteome Software | https://www.proteomesoftware.com/products/scaffold-5 |
| PyMOL (v2.4.1) | Schrodinger | https://pymol.org/ |
| Web 3DNA 2.0 | Zheng et al.<sup>85</sup> | http://web.x3dna.org/ |
| MolProbity (v4.2) | Chen et al.<sup>86</sup> | http://molprobity.biochem.duke.edu/ |
| PHENIX (v1.20) | Terwilliger et al.<sup>81</sup> | https://phenix-online.org/ |
| Code used for data analysis and graph plotting | This paper | DOI: 10.5281/zenodo.11529109 |
| Other |  |  |
| Amersham Protran, 0.45 µm pores | Cytiva | Cat#10600062 |
| Amersham Hybond-XL | Fisher Scientific | Cat#45001147 |

### RESOURCE AVAILABILITY

#### Lead contact

Further information and requests for resources and reagents should be directed to and will be fulfilled by the lead contact, David P. Bartel.

#### Materials availability

Plasmids generated in this study are deposited at Addgene.

#### Data and code availability

- Gel images, microscopy images, luciferase readings, mass spectrometry data, and reaction assay data have been deposited at Mendeley Data. Accession numbers are listed in the key resources table. These data are publicly available as of the date of publication.
- All original code used for data analysis and graphing has been deposited at Zenodo and is publicly available as of the date of publication. DOIs are listed in the key resources table.
- Any additional information required to reanalyze the data reported in this paper is available from the lead contact upon request.

### EXPERIMENTAL MODEL AND STUDY PARTICIPANT DETAILS

#### Cell lines

HEK293FT, HEK293T, and HeLa cells were cultured in DMEM (VWR, 45000-304) supplemented with 10% FBS (Takara Bio, 631367), at 37 °C in 5% CO_2_. Cells were passaged 1:10 every 3−4 days.

### METHOD DETAILS

#### In vitro transcription of target RNAs

Single-stranded DNA templates (Table S2), each containing a T7 promoter sequence, were synthesized (IDT), purified on a 12% or 15% urea-polyacrylamide gel, and resuspended in water. Each template DNA was annealed with an oligonucleotide complementary to the T7 promoter sequence (Table S2), and then used in an in vitro transcription reaction containing 0.5 µM annealed DNA template, 5 mM ATP, 2 mM UTP, 5 mM CTP, 8 mM GTP, 5 mM DTT, 40 mM Tris pH 7.9, 2.5 mM spermidine, 26 mM MgCl_2_, 0.01% (v./v.) Triton X-100, 5 mM DTT, SUPERase•In (1 U/µL, Invitrogen, AM2694), thermostable Inorganic Pyrophosphatase (0.0083 U/µL, New England Biolabs, M0296), and T7 RNA Polymerase (purified in-house). The reaction was incubated at 37°C for 3−4 h, before RQ1 DNase (0.037 U/µL, Promega, M6101) was added, followed by 30 min of incubation at 37°C. The final reaction mix was desalted using Micro Bio-Spin P-30 columns (Bio-Rad, 7326250), then denatured by incubating with 0.5X volume of gel-loading buffer (8 M urea, 25 mM EDTA, 0.025% [w./v.] xylene cyan, 0.025% [w./v.] bromophenol blue) or Gel Loading Buffer II (Invitrogen, AM8547) at 90°C for 1 min. The full-length RNA product was then purified on a 12% or 15% urea-polyacrylamide gel and resuspended in water.

#### Preparation of radiolabeled RNAs

Guide RNAs (Table S2) were synthesized with a 5′ monophosphate (IDT), purified on a 15% urea-polyacrylamide gel, and then resuspended in water. To radiolabel, guide RNAs were first dephosphorylated using shrimp alkaline phosphatase (rSAP, 0.05 U/µL; New England Biolabs, M0371) at 37 °C for 30 min, followed by heat-inactivation at 75°C for 5 min and desalting using Micro Bio-Spin P-6 columns (Bio-Rad, 7326221). Dephosphorylated guide RNAs were then radiolabeled using T4 Polynucleotide Kinase (0.33 U/µL, New England Biolabs, M0201) and 0.42 µM [γ-^32^P] ATP (PerkinElmer, NEG035C001MC) at 37°C for 1.5 h, before a brief chase with 0.16 mM ATP at 37°C for 15 min, followed by desalting using Micro Bio-Spin P-6 columns and denaturation with gel-loading buffer or Gel Loading Buffer II. Radiolabeled guide RNAs were purified on a 15% urea-polyacrylamide gel, and then resuspended in water. Guide RNAs intended for purification of RISCs for hydroxyl footprinting were labeled similarly with up to 5 µM [γ-_32_P] ATP, without chase with cold ATP.

Short target RNAs, containing a seed (7mer-m8 or 8mer)-only binding site for titration quantification (Table S2), were synthesized (IDT), purified on a 15% urea-polyacrylamide gel, and then resuspended in water. These RNAs were then radiolabeled with T4 Polynucleotide Kinase and 2.5 µM [γ-^32^P] ATP at 37°C for 1.5 h, followed by the chase and purification described for radiolabeled guide RNAs.

In vitro-transcribed target RNAs (Table S2) were radiolabeled with ^32^P at either a 5′ monophosphate or a 7-methylguanylate cap. The identity of target 5′ chemistry had no detectable impact on our biochemical assays. When labeling at a 5′ monophosphate, RNAs were first dephosphorylated using Quick CIP (0.1 U/µL, New England Biolabs, M0525) at 37°C for 15 min, followed by heat-inactivation at 80°C for 3 min. Dephosphorylated RNAs were then radiolabeled using T4 Polynucleotide Kinase (0.67 U/µL) and 0.84 µM [γ-^32^P] ATP, in CutSmart buffer (New England Biolabs) supplemented with 5 mM DTT, at 37°C for 1.5 h, before a brief chase with 0.32 mM ATP at 37°C for 15 min. When labeling at a 7-methylguanylate cap, RNAs were capped using the *Vaccinia* capping system (New England Biolabs, M2080) with [α-^32^P] GTP (PerkinElmer, NEG006H250UC), according to manufacturer’s instructions. Reactions from either labeling method were desalted using Micro Bio-Spin P-30 columns, denatured with gel-loading buffer or Gel Loading Buffer II, purified on a 12% or 15% urea-polyacrylamide gel, and then resuspended in water.

#### Purification of RISCs

5′-phosphorylated guide and passenger RNAs (Table S2) were synthesized (IDT), purified on a 15% urea-polyacrylamide gel, and then resuspended in water. Passenger RNAs were designed to create mismatches in the seed and 2-nt 3′-overhangs on both ends of the guide−passenger RNA duplexes, in order to direct the correct strand selection by AGO2.^87^ Guide−passenger RNA duplexes were assembled using 1 µM each of guide and passenger RNAs, with 1−10% ^32^P-radiolabeled guide RNA, in annealing buffer (30 mM Tris-HCl pH 7.5, 100 mM NaCl, and 1 mM EDTA). RNAs were annealed by heating to 90°C and then slowly cooling over 1.5 h to 30°C, before being chilled on ice and stored at −20°C.

Each RISC was assembled in lysate containing tagged AGO2 protein and then purified by oligo-affinity capture and competitive elution.^50^ Lysates containing AGO2 proteins, N-terminally tagged with a 3xFLAG sequence, were generated as follows: HEK293FT cells were co-transfected at ∼80% confluency with the pcDNA3.3−3xFLAG−AGO2^49^ (Addgene plasmid #136687) and pMaxGFP (Lonza) plasmids at 4:1 ratio (by mass) using Lipofectamine 2000 (Invitrogen, 11668019) in Opti-MEM (Gibco, 51985091), according to manufacturers’ instructions. After 24 h, cells were harvested and lysed in hypotonic buffer (10 mM HEPES pH 7.4, 10 mM KOAc, 1.5 mM Mg(OAc)_2_, 0.5 mM DTT, with cOmplete Mini EDTA-free Protease Inhibitor Cocktail [1 tablet per 10 mL, Roche, 11836170001]) by passing through a 23G × 1 inch needle 30 times. Cell lysate was clarified by centrifugation at 2,000*g* at 4°C for 10 min four times, then once for 30 min. The supernatant of this hypotonic lysate was reequilibrated by adding 11% volume of re-equilibration buffer (300 mM HEPES pH 7.4, 1.4 M KOAc, 30 mM Mg(OAc)_2_, with 1 tablet per 10 mL cOmplete Mini EDTA-free Protease Inhibitor Cocktail), before further clarification by centrifugation at 100,000*g* at 4°C for 20 min. 20% lysate-volume of 80% (v./v.) glycerol was added before flash-freezing in liquid nitrogen in aliquots of 600 µL for storage at −150°C.

To assemble each RISC, 300−600 µL of cell lysate was incubated for 2 h at 25°C with 11% lysate-volume of assembled guide−passenger duplex, with the final concentration of the duplex at 100 nM. Loaded lysate was centrifuged at 21,000*g* at 4°C for 10 min, and the supernatant was incubated with 150 µL of a slurry of Dynabeads MyOne Streptavidin C1 (Invitrogen, 65002) pre-bound to 75 pmol of capture oligonucleotide (Table S2), at 25°C with shaking at 1,300 rpm for 1.25 h. These pre-bound streptavidin beads were prepared by equilibrating the beads according to manufacturer’s instructions, followed by incubation with 75 pmol of 3′-end biotinylated, fully 2′-*O*-methylated oligonucleotides (IDT) containing a seed-only 8mer-site for the guide RNA (Table S2), at 25°C with shaking at 1,300 rpm for 30 min. The beads were then washed in 300 µL equilibration buffer (18 mM HEPES pH 7.4, 100 mM KOAc, 1 mM Mg(OAc)_2_, 0.01% [v./v.] IGEPAL CA-630 [Sigma-Aldrich, I3021], 0.1 mg/mL BSA [New England Biolabs, B9000], and 0.01 mg/mL yeast tRNA [Life Technologies, AM7119]) before use. After incubation with the loaded lysate, the beads were washed five times with 200 µL equilibration buffer, and five times with 200 µL capture-wash buffer (18 mM HEPES pH 7.4, 2 M KOAc, 1 mM Mg(OAc)_2_, 0.01% [v./v.] IGEPAL CA-630, 0.1 mg/mL BSA, and 0.01 mg/mL yeast tRNA). Washed beads were incubated with 150 µL of competitor elution solution (0.75 µM of 3′-biotinylated DNA competitor oligonucleotides [IDT] complementary to the capture oligonucleotides (Table S2), 18 mM HEPES pH 7.4, 1 M KOAc, 1 mM Mg(OAc)_2_, 0.01% [v./v.] IGEPAL CA-630, 0.1 mg/mL BSA, and 0.01 mg/mL yeast tRNA) at 25 °C with shaking at 1,300 rpm for 2 h. Purifications of AGO2−miR-451a and AGO2–miR-451a.M1 complexes used competitor oligonucleotides without 3′-biotinylation, with negligible difference in performance. The eluate was then incubated with 20 µL of an anti-FLAG M2 magnetic beads slurry (Millipore, M8823), pre-equilibrated in equilibration buffer according to manufacturer’s instructions, at 25°C with shaking at 1,100 rpm for 2 h. The beads were washed three times with 200 µL of equilibration buffer, and three times with 500 µL of equilibration buffer, before incubation in 60 µL 3xFLAG elution solution (146 ng/µL of 3xFLAG peptide [Sigma-Aldrich, F4799] in 0.97X equilibration buffer) at 25°C with shaking at 1,100 rpm for 1 h. The eluate was supplemented with glycerol and DTT to a final storage buffer condition of 13.0 mM HEPES pH 7.4, 72.3 mM KOAc, 0.723 mM Mg(OAc)_2_, 5 mM DTT, 0.0723 mg/mL BSA, 0.00723 mg/mL yeast tRNA, 0.00723% (v./v.) IGEPAL CA-630, and 20% (v./v.) glycerol, before flash-freezing in liquid nitrogen for storage at −80°C.

To produce complexes with mutant AGO2 proteins, including AGO2^D669A^, AGO2^S387A^, and AGO2^S387D^, the corresponding plasmids were generated by site-directed mutagenesis of the pcDNA3.3−3xFLAG−AGO2 plasmid, by PCR (Table S2) using KAPA HiFi HotStart Ready Mix (Roche, 07958935001). Plasmid sequences were confirmed by Sanger sequencing.

AGO2^D669A^−guide complexes used for hydroxyl-radical footprinting were purified in a similar manner but without inclusion of glycerol in the final storage buffer (17.4 mM HEPES pH 7.4, 96.6 mM KOAc, 0.966 mM Mg(OAc)_2_, 5 mM DTT, 0.0966 mg/mL BSA, 0.00966 mg/mL yeast tRNA, and 0.00966% [v./v.] IGEPAL CA-630). Small aliquots were flash-frozen in liquid nitrogen for storage at −80 °C.

#### Quantification of purified RISCs

Each purified RISC was first run on a 15% urea-polyacrylamide gel alongside the guide−passenger duplex used for its purification. Gels were frozen at −20 °C while exposing a phosphorimager plate, and radioactivity was then imaged and quantified. Radioactivity of the guide in the purified RISC relative to that in the duplex (for which the concentration was known) provided an estimate for the concentration of the purified complex.

Concentrations of purified complexes were then more accurately determined using a titration assay. A dilution series of RISC concentrations that were designed to not exceed 1 nM, were each incubated with 1 nM of radiolabeled oligonucleotides harboring a single canonical 7mer-m8 or 8mer binding site for the corresponding guide RNA. Binding reactions were incubated at 37°C for 1 h, in a final binding reaction buffer of 15.5 mM HEPES pH 7.4, 86.2 mM KOAc, 0.862 mM Mg(OAc)_2_, 5 mM DTT, 0.05 mg/mL BSA, 0.005 mg/mL yeast tRNA, and 10% (v./v.) glycerol. To carry out filter-binding experiments, nitrocellulose (Amersham Protran, 0.45 µm pores; Cytiva, 10600062) and nylon (Amersham Hybond-XL; Fisher Scientific, 45001147) membrane filters were cut into discs of 0.5-inch diameter and equilibrated at 25°C for at least 20 min in filter-binding buffer (18 mM HEPES pH 7.4, 100 mM KOAc, 1 mM Mg(OAc)_2_). Each nitrocellulose disc was stacked on top of a nylon disc, then placed on a circular pedestal mounted on a Visiprep SPE Vacuum Manifold (Supelco, 57250-U), set at approximately −20 kPa. 10 µL of reaction was applied to stacked filter membrane discs, followed by 100 µL of ice-cold filter-binding wash buffer (18 mM HEPES pH 7.4, 100 mM KOAc, 1 mM Mg(OAc)_2_, 5 mM DTT). Filter membrane discs were separated, airdried, then imaged by phosphorimaging and quantified.

Fractions of target RNA bound across reactions were used to quantify stock concentration of RISCs, assuming a 1:1 stoichiometric ratio of RISC and bound target and *K*_D_ of the binding reaction far below the total target concentration. Datasets that did not approach target saturation (≤ 60% bound) were fit by linear regression in R (lm) to the following equation:

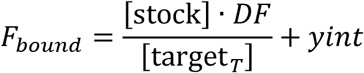

where *F*_bound_ represents fraction of target bound, [target_T_] represents concentration of total target oligonucleotide (1 nM), [stock] represents stock concentration of RISC, *DF* represents the dilution factor, and *yint* represents the *y*-intercept, which was near zero for all datasets.

Datasets approaching saturation were instead fit to a quadratic equation by nonlinear least-squares regression in R using the Levenberg-Marquardt algorithm (nlsLM from the R package minpack.lm):

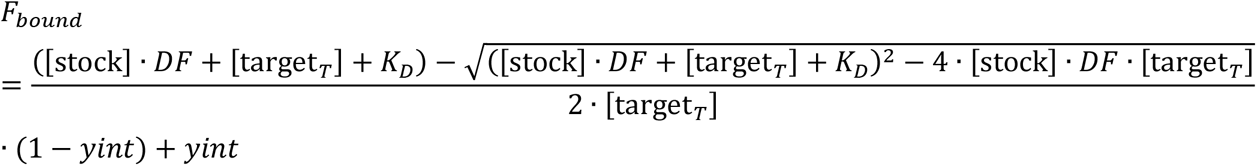

where *K*_D_ additionally represents the dissociation constant for the affinity between RISC and the seed-match target. [stock] was initialized at 30,000 pM and limited to the numerical range (0, 10^6^); *yint* was initialized at 0.1 and limited to the numerical range (0, 1); *K*_D_ was initialized at 10^2^ pM, limited to the numerical range (10^−2^, 10^4^), and fit in log-transformed space. For lsy-6, the nonlinear least-squares regression did not produce a confident estimation of stock concentration (*p* > 0.05), and thus the results from a linear regression of data points before saturation were used instead.

#### In vitro slicing assays

Target RNAs for slicing assays were designed with a perfectly or partially complementary slicing site, flanked by poly(AAC) or poly(UUC) sequences designed to reduce secondary structure. Single-turnover assays were conducted with 0.02−0.10 nM of radiolabeled target RNA and a dilution series of RISC concentrations (at 0.09−33 nM and 4.5−660-fold excess over the target across all assays). Target RNA was preincubated in slicing-reaction buffer (18 mM HEPES pH 7.4, 100 mM KOAc, 1 mM Mg(OAc)_2_, 5 mM DTT, 0.01% [v./v.] IGEPAL CA-630) at 37°C for 1.5 min, while the RISC was preincubated in storage buffer at 37°C for 0.5 min, before an equal volume of preincubated RISC was added to preincubated target RNA in buffer, to a final reaction condition of 15.5 mM HEPES pH 7.4, 86.2 mM KOAc, 0.862 mM Mg(OAc)_2_, 5 mM DTT, 0.05 mg/mL BSA, 0.005 mg/mL yeast tRNA, 0.01% (v./v.) IGEPAL CA-630, and 10% (v./v.) glycerol. Reactions were incubated at 37°C. At select time points, aliquots of the reaction were quenched by mixing rapidly with gel-loading buffer or Gel Loading Buffer II on ice. Quenched reaction aliquots were denatured at 90°C for 1 min, and then run on a 15% urea-polyacrylamide gel. Gels were frozen at −20°C while exposing a phosphorimager plate, and radioactivity was imaged and then quantified.

Multiple-turnover assays were conducted with 4 nM radiolabeled target and ∼0.25 nM RISC. Conditions for preincubation, reaction, and quenching, and methods of phosphorimaging and quantification, were otherwise the same as those of single-turnover assays.

#### Model-fitting of single-turnover slicing kinetics

Guesses were first determined for each dataset. *k*_on_ was initialized at 0.0167 nM^−1^ s^−1^ for datasets with substantial contribution from slow association kinetics, and 0.167 nM^−1^ s^−1^ for datasets without. Datasets were defined to have substantial contribution from slow association kinetics if deviation in mean fraction of target sliced across different RISC concentrations was more than 0.15 at any time point. *k*_slice_ was initialized at 0.0167 s^−1^, *k*_phase2_ at 3.33 × 10^−6^ s^−1^, and *F*_a_ at 0.95 × the highest fraction of target sliced measured in the dataset. These were used to generate guesses by nonlinear least-squares regression in R to an approximation equation. Datasets were fit using the Levenberg-Marquardt algorithm (nlsLM), for no more than 50 iterations, to the following equation, assuming pseudo-steady-state of enzyme-substrate complex concentration:

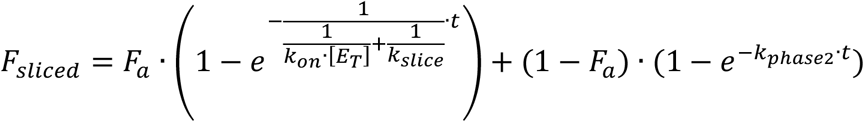

where *F*_sliced_ represents fraction of target sliced, *t* represents time in s, *k*_on_ represents the association rate constant in nM^−1^ s^−1^, [E_T_] represents the total concentration of RISC in nM, *k*_slice_ represents the slicing rate constant in s^−1^, *k*_phase2_ represents the slow second-phase slicing rate constant in s^−1^, and *F*_a_ represents the height of the first phase. *k*_on_ was limited to the numerical range (10^−6^, 1), where 1 nM^−1^ s^−1^ was the diffusion limit. *k*_slice_ was limited to the numerical range (10^−6^, 1). *k*_phase2_ was limited to the numerical range (1.67 × 10^−6^, 1.67 × 10^−4^), where 1.67 × 10^−6^ s^−1^ was the approximate detection limit based on the longest time points tested. *F*_a_ was limited to the numerical range (0.3, 0.999). If the nonlinear least-squares regression failed, the initial values were used with *F*_a_ adjusted to 0.85. For datasets with slow association kinetics, the fitted value of *F*_a_ was adjusted by a factor of 0.9 to account for overestimation of the plateau by the pseudo-steady-state approximation model.

These guesses were then used as initial values to fit the dataset to an ODE model to calculate accurate values:

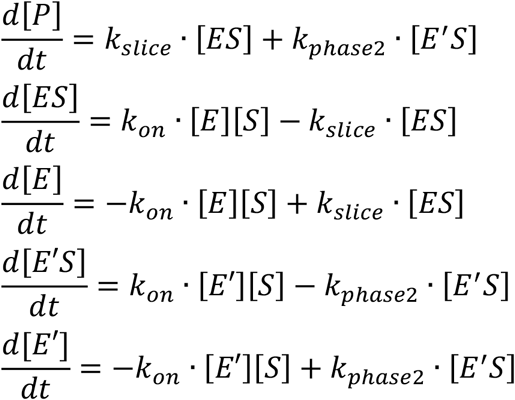

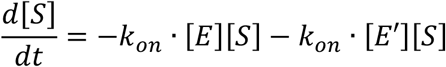

With

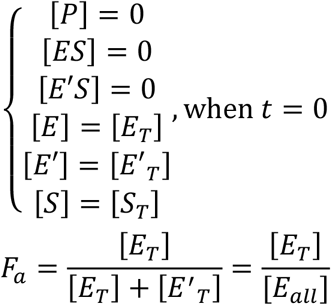

such that, at each time point,

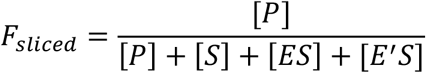

where *F*_sliced_ represents fraction of target sliced, *t* represents time in s, [E_T_] represents total concentration of functionally intact RISC in nM, [E’_T_] represents total concentration of defective RISC in nM, [E_all_] represents total concentration of all RISCs, *F*_a_ represents the fraction of RISCs that is functionally intact, *k*_on_ represents the association rate constant in nM^−1^ s^−1^, *k*_slice_ represents the slicing rate constant in s^−1^, and *k*_phase2_ represents the defective slicing rate constant in s^−1^.

Other biphasic ODE equations were also attempted to model the biphasic behavior as each of the following: (1) a small fraction of inactive RISCs binding to target RNAs but unable to slice, instead dissociating at *k*_phase2_; (2) a small fraction of the enzyme-substrate complex adopting a slicing-incompetent conformation upon binding, dissociating at *k*_phase2_; (3) a small fraction of the enzyme-substrate complex adopting a slicing-incompetent conformation upon binding, changing to a slicing-competent conformation at *k*_phase2_; (4) a small fraction of the enzyme-substrate complex adopting a slicing-defective conformation upon binding, slicing at *k*_phase2_; and (5) a small fraction of the enzyme-substrate complex undergoing unloading of the guide, leaving inactive naked guide RNA blocking the target RNA, dissociating at *k*_phase2_. Although the fits for these rejected models were indistinguishable from those for the accepted model, these models were rejected in favor of the accepted model based on orthogonal experimental results (Figures S2D, S2F, and S5D). Importantly, the choice of models did not change any of our other interpretations or conclusions.

Fitting to the ODE model was carried out using the modFit function, with the Levenberg-Marquardt algorithm, from the FME package in R,^83^ for up to 200 iterations. At each iteration, reaction time curves were calculated for each pair of RISC and target concentration values using the Livermore solver for ordinary differential equations (LSODE) from the deSolve package (ode function) in R.^82^ Calculations were conducted using the backward differentiation formula with a full Jacobian matrix internally calculated by lsode (option mf = 22). Relative tolerance was set to 10^−6^, while absolute tolerance was set to 10^−10^. At each iteration, the cost function was defined to be the absolute deviation in fraction of target sliced between empirical and ODE-predicted values. Kinetic constants were fit in log-transformed space. *k*_on_ was limited to the numerical range (0, 1), where 1 nM^−1^ s^−1^ was the diffusion limit. *k*_slice_ was limited to the numerical range (0, ∞). *k*_phase2_ was limited to the numerical range (1.67 × 10^−6^, 3.33 × 10^−3^). *F*_a_ was limited to the numerical range (0.3, 1.0). If *k*_on_ could not be confidently fit for fast-binding reactions due to trivial contribution from binding kinetics, model fitting was repeated with *k*_on_ constrained to the diffusion limit at 1 nM^−1^ s^−1^. If *k*_phase2_ could not be confidently fit due to insufficiently long time points to resolve the second phase, model fitting was repeated with *k*_phase2_ constrained to zero.

As RISC was in excess over the target, exact initial concentrations of target RNA had negligible impact on the reaction rates. While ODE model fitting took into account exact initial concentrations of target RNA, reaction curves in figures were plotted with the initial target concentration at 0.02 nM for simplicity.

For some guides with rapid binding kinetics that did not contribute meaningfully to the reaction kinetics, reaction assays using other purifications of RISC as biological replicates were conducted at a single RISC concentration, and were fit with the binding kinetics constrained to the diffusion limit (1 nM^−1^ s^−1^).

For some assays with mismatched targets (miR-124.M2, 16-bp target; miR-196a, 16-bp and mm17GA targets; miR-196a.M1, 16-bp and mm17AA targets), slicing kinetics were too slow for the plateau to be resolved even at very long time points, with no detectable contribution from association kinetics. Therefore, these datasets were instead fit by nonlinear least-squares regression in R using the Levenberg-Marquardt algorithm (nlsLM), for no more than 100 iterations, to the following equation:

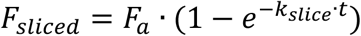

where *F*_a_ was set to the fitted value from the same guide with its perfectly complementary target. *k*_slice_ was initialized at 1.67 × 10^−4^ s^−1^, limited to the numerical range (0, ∞), and fit in log-transformed space.

#### Model-fitting of multiple-turnover slicing kinetics

Datasets were fit to the multiple-turnover ODE model, accounting for product release and re-binding:

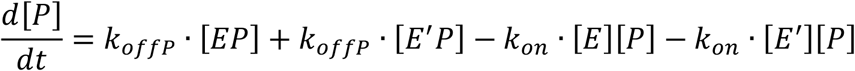

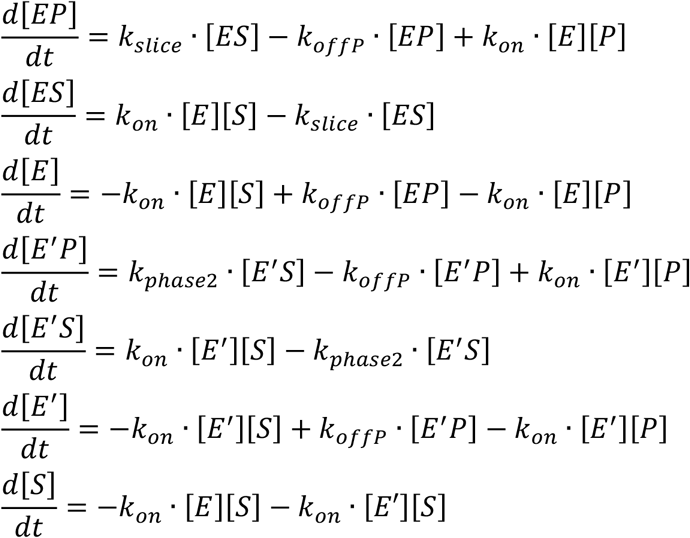

With

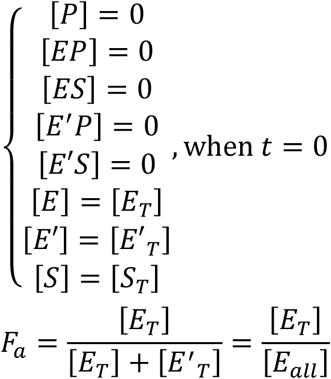

such that, at each time point,

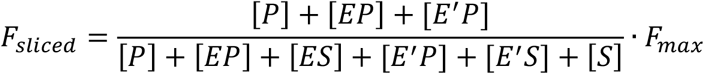

where, in addition to parameters defined in the single-turnover ODE equations, *F*_max_ represents the maximum plateau height of the fraction of target sliced, and *k*_offP_ represents the rate constant of product release in s^−1^. Values of *k*_on_, *k*_slice_, *k*_phase2_, and *F*_a_ were taken from single-turnover slicing assay results of the same RISC and target. *k*_offP_, *F*_max_, and [E_all_] were fit to the model, with *k*_offP_ being fit in log-transformed space, and [E_all_] being fit as a deviation from the experimentally intended concentration. For datasets with at least one measured fraction of target sliced exceeding 0.2 due to rapid catalytic kinetics, *k*_offP_ was initialized at 0.167 s^−1^ and limited to the numerical range (0, ∞), *F*_max_ was initialized at 0.95 and limited to the numerical range (0, 1), and [E_all_] was initialized at the experimentally intended concentration and limited to two-fold above or below this value. Fitted values of *F*_max_ were between 0.94 and 0.95 across five datasets.

For datasets with all values of fraction of target sliced below 0.2, *k*_offP_ was initialized instead at 0.0167 s^−1^, and *F*_max_ was not fit and instead constrained to 1.

#### Filter binding for RISC binding kinetics

Filter binding was conducted with target RNAs bearing a mismatch at position 10, or mismatches at positions 10−11 for lsy-6, to best mimic the binding of a perfectly complementary target without slicing. Filter membrane discs were prepared and preincubated in the same way as in filter-binding assays for RISC quantification. Target RNA was preincubated in slicing-reaction buffer at 37°C for 1.5 min, while the RISC was preincubated in storage buffer at 37°C for 0.5 min, before an equal volume of RISC was added to the preincubated target RNA in buffer and mixed, with the final concentration of RISC at 0.5 or 1 nM, and the final concentration of target at 10-fold below that of RISC. Reactions were incubated at 37°C. At select time points, 5 µL of binding reaction was applied to stacked filter membrane discs, followed by 100 µL of ice-cold filter-binding wash buffer. The filter membrane discs were then quantified in the same way as in titration filter-binding assays for RISC quantification.

The fractions of target bound across time points were then fit to an exponential equation by nonlinear least-squares regression in R using the Levenberg-Marquardt algorithm (nlsLM), for no more than 100 iterations:

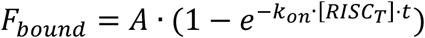

where *F*_bound_ represents the fraction of target bound, *A* represents the maximum fraction of target bound at the plateau, *k*_on_ represents the association rate constant in nM^−1^ s^−1^, [RISC_T_] represents the total RISC concentration in nM, and *t* represents time in s. This equation approximates association rate *k*_on_[RISC] to be *k*_on_[RISC_T_] and dissociation to be negligible, as the concentration of RISC was in vast excess over both the target concentration and *K*_D_ of binding. *A* was initialized at 0.8 and limited to the range (0, 1), and *k*_on_ was initialized at 0.0333 nM^−1^ s^−1^ and limited to the numerical range (0, 1), where 1 nM^−1^ s^−1^ was the diffusion limit. For lsy-6, *k*_on_ was too fast to resolve even with the first time point measured, and was therefore ≥ 1 nM^−1^ s^−1^ (≥ 60 nM^−1^ min^−1^).

#### Filter binding of slicing reactions

Slicing reactions were prepared and incubated similarly as described. Instead of quenching the reactions, they were filtered through a single pre-equilibrated nitrocellulose membrane. Individual discs of nitrocellulose membrane filters were placed on a circular pedestal mounted on a vacuum manifold, set at approximately −20 kPa. 5 µL of reaction was applied to each filter membrane disc, immediately followed by 100 µL of ice-cold filter-binding wash buffer. The filtrate was collected and mixed with 11 µL of 3 M NaCl, 1 µL of 25 mg/mL yeast tRNA, 5 µL of GlycoBlue co-precipitant (Invitrogen, AM9516), and 320 µL of ethanol, and incubated at −80°C for at least 1 h. The nitrocellulose membrane was air-dried briefly, then incubated in 420 µL of Proteinase K elution solution (47.3 mM Tris-HCl pH 7.5, 47.3 mM NaCl, 9.46 mM EDTA, 0.946% [v./v.] SDS, 0.01 mg/mL yeast tRNA, 1 mg/mL Proteinase K [Invitrogen, 25530049]) at 65°C with shaking at 1,200 rpm for 45 min. RNA was isolated from the proteinase digestion eluate by extraction with phenol:chloroform:isoamyl alcohol (25:24:1) (Sigma-Aldrich, P2069) and chloroform. The resultant aqueous phase was mixed with 40 µL of 3 M NaOAc, 2 µL of 2.5 mg/mL yeast tRNA, 5 µL of GlycoBlue co-precipitant, and 1000 µL of ethanol, and incubated at −80°C for at least 1 h. RNA species from the filtrate and membrane eluate were precipitated by centrifugation at 4°C, and dried RNA pellets were each resuspended in 15 µL of a 1:1 (by volume) mixture of Gel Loading Dye II and filter-binding buffer. Samples were denatured by incubating at 90°C with shaking at 1,500 rpm for 5 min, and run on a 15% urea-polyacrylamide gel. Gels were imaged and quantified as described.

**Thermodynamic modeling of slicing kinetics for 16-bp targets**

The relationship between the relative *k*_slice_ for slicing 16-bp targets compared to that for slicing perfect targets (*k*_slice (16-bp)_ / *k*_slice (perfect)_) and the predicted base-pairing energy at positions 9−12 was modeled as the following equation, where energy gained from base-pairing is used to overcome an energetic barrier to slicing, assuming the slicing kinetics of perfectly complementary targets correspond to the maximal rates:

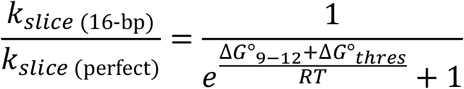

where Δ*G*°_9−12_ represents the predicted base-pairing energy at positions 9−12, Δ*G*°_thres_ represents the energetic threshold to be overcome by Δ*G*°_9−12_, *R* represents the molar gas constant, and *T* represents the temperature at 310.15 K (37°C). Data were fit to this equation in log-scale using nonlinear least-squares regression in R, using the Levenberg-Marquardt algorithm (nlsLM). Δ*G*°_thres_ was initialized at 10 kcal mol^−1^ and not bounded.

#### Hydroxyl-radical footprinting of AGO2^D669A^−RNA complexes

Base-hydrolysis ladders were generated by incubating 30 nM of radiolabeled guide RNAs with 50 mM Na_2_CO_3_, in a total volume of 10 µL, at 90°C for 4 min, followed by quenching with 20 µL of ice-cold 90% (v./v.) gel-loading buffer and 100 mM Tris-HCl acid. RNase T1 ladders were generated by first preincubating 3 nM of radiolabeled guide RNAs with 5 mM EDTA, 0.75 mg/mL GlycoBlue, 0.025 mg/mL yeast tRNA, and 0.88X RNA Sequencing Buffer, in a total volume of 100 µL, at 50°C for 5 min, then at 25°C for 3 min. 1 µL of RNase T1 (Invitrogen, AM2283) was added and mixed, and the reaction incubated at 25°C for 20 min, before adding 10 µL of 1 M DTT and extracting the RNA using phenol:chloroform:isoamyl alcohol (25:24:1) and chloroform. The aqueous phase was mixed with 200 µL of the Precipitation/Inactivation Buffer provided with RNase T1, and incubated at −80°C for 1 h. After centrifugation, the precipitate was resuspended in 30 µL of Gel Loading Dye II and denatured at 90°C for 2 min.

To provide a normalization standard for maximum hydroxyl-radical reactivity, radiolabeled guide RNAs were purified from AGO2^D669A^−guide complexes by phenol:chloroform:isoamyl alcohol (25:24:1) and precipitation. AGO2^D669A^−guide complexes or naked guide RNAs were preincubated in storage buffer without glycerol at 37°C for 30 s or 2 min. Non-radiolabeled target RNAs were preincubated in slicing-reaction buffer at 37°C for 30 s or 2 min. AGO2^D669A^−guide complexes were mixed with target RNAs in buffer to a total volume of 15 µL, with the final concentration of AGO2^D669A^−guide at 2−4 nM and target at 4−6-fold excess. The binding reaction mix was incubated at 37°C for indicated lengths of time. Meanwhile, Fenton’s reagent was prepared in buffer on ice as two separate solutions for each reaction: Solution A, containing 14.7 mM EDTA, 13.3 mM ammonium iron(II) sulfate (Sigma-Aldrich, 09719), and 33.3 mM DTT; and Solution B, containing 1% (v./v.) hydrogen peroxide (Sigma-Aldrich, 216763), 45 mM HEPES pH 7.4, 250 mM KOAc, 4.5 mM Mg(OAc)_2_, and 0.025% (v./v.) IGEPAL CA-630. Solutions A and B were preincubated separately at 37 °C for 20 s. To initiate footprinting, the binding reaction was first rapidly mixed with 3 µL of Solution B, and then immediately mixed with 2 µL of Solution A, to a final total volume of 20 µL. The final footprinting reaction condition contained 1.5−3 nM AGO2^D669A^−guide or naked guide RNA, 4−6-fold excess target RNA, 18 mM HEPES pH 7.4, 100 mM KOAc, 1.2 mM Mg(OAc)_2_, 0.015 mg/mL BSA, 0.0015 mg/mL yeast tRNA, 0.01 (v./v.) IGEPAL CA-630, 8.75 mM DTT, 2.2 mM EDTA, 2 mM ammonium iron(II) sulfate, and 0.1% (v./v.) (29.5 mM) hydrogen peroxide. The footprinting reaction mix was incubated at 37°C for 2 min, before quenching with 5 µL of ice-cold 0.5 M sodium sulfite (Sigma-Aldrich, S0505) and incubating on ice for at least 5 min. To provide a normalization standard for minimum reactivity, a quenched reaction was generated in which the incubated binding reaction was mixed with a pre-quenched mix of Solutions A and B and sodium sulfite on ice. 25 µL of each quenched reaction mix was added to 455 µL of ice-cold 0.3 M NaCl, and RNA species were extracted by phenol:chloroform:isoamyl alcohol (25:24:1) and chloroform. The aqueous phase was mixed with 5 µL of 5 mg/mL linear acrylamide co-precipitant (Invitrogen, AM9520) and 1000 µL of ethanol for precipitation. Dried pellets were resuspended in 10 µL of 200 mM Tris-HCl acid, 75% (v./v.) Gel Loading Dye II, and 25−50 nM non-radiolabeled guide RNA. The excess non-radiolabeled guide RNA was included so that it would anneal with complementary target RNA in the mix, thereby displacing radiolabeled guide-RNA species that would otherwise be paired to target RNA. Samples were denatured at 60°C for 10 min, and then resolved on a 20% urea-polyacrylamide sequencing gel, alongside previously prepared radiolabeled ladders of the guide RNAs. Gels were imaged and quantified as described.

Reactivity values were normalized by linearly rescaling the value at each nucleotide position relative to the mean reactivities of quenched-reagent samples (set to 0) and naked-guide samples (set to 1). Non-specific degradation band(s) introduced when preparing naked-guide samples for one replicate each of miR-196a and miR-124.M2 were identified by gel electrophoresis and corrected by replacing the corresponding experimental values with the mean of values at neighboring positions (Figure S6B, asterisk). One replicate for miR-430a bound to a seed+supplementary pairing target was discarded due to denatured complex, as indicated by a uniform and substantially elevated reactivity across all nucleotide positions, half of which were > 2 standard deviations above the mean across replicates.

#### Mass spectrometry of purified RISCs

10 ng of AGO2 protein from each RISC, as quantified by titration filter binding, was denatured with 4X LDS sample buffer (Invitrogen, NP0007) and run on an SDS-polyacrylamide gel (NuPAGE 4 to 12%, Bis-Tris [Invitrogen, NP0322]) in MOPS buffer (Invitrogen, NP0001), according to manufacturer’s instructions. Gels were stained with Imperial Blue Stain (Thermo Scientific, 24615). Bands corresponding to AGO2 were excised and cut into pieces, then stored in 50% (v./v.) methanol. Proteins were eluted, digested with trypsin, and run on the Thermo Scientific Orbitrap Elite mass spectrometer with a Waters NanoAcuity UPLC system. Alignment of peptides and identification of modifications were carried out using either the Proteome Discoverer (Thermo Fisher) or Mascot (Matrix Science) software, followed by Scaffold 5 (Proteome Software). The label-free total ion chromatogram (TIC) signal for each assigned peak was used for relative quantitative analyses. Fraction of phosphorylation at S387 was estimated based on quantification of the “SAS^387^FNTDPYVR” peptide observed.

#### Luciferase reporter assays for RNAi-mediated knockdown

Reporter plasmids each featured a pUC origin of replication, an ampicillin resistance gene, and two minimal CMV promoters in divergent directions, with a common enhancer element in between. One promoter expressed a firefly luciferase that was used for reporter normalization. The other promoter expressed the reporter construct that contained the slicing site. The cap-dependent translation of its first ORF, which harbored the slicing site, was regulated by an IRE in the mRNA 5′ UTR. The 22-nt-long perfectly complementary slicing site for each guide RNA was introduced in the first ORF as part of a 24-nt-long region that linked the two ORF regions, preserving the codon frame. For each guide and ORF context, the frame of the slicing site was chosen to avoid any stop codons, rare codons, or highly hydrophobic amino acid residues. To measure the abundance of the reporter transcript independently from the regulation by the IRE, a second ORF was placed after a downstream EMCV IRES. It expressed a NanoLuc luciferase with a C-terminal PEST degradation sequence from mouse ornithine decarboxylase.^88^ The PEST degron destabilized the NanoLuc protein, enabling faster approach to steady-state reporting of the mRNA abundance after treatments. The entire reporter construct transcript was approximately 2 kb in length. Both mRNAs terminated with the SV40 polyadenylation-signal sequence.

Two reporter template plasmids without slicing sites were each constructed by Gibson assembly from six fragments, including those encoding 3xFLAG or 3xHA, sfGFP or SUMO^Eu1^, EMCV IRES, NanoLuc, PEST degron, and a plasmid backbone containing the remaining elements, mixed in 5:2:2:2:2:1 ratio by molarity, using the NEBuilder HiFi DNA Assembly Master Mix (New England Biolabs, E2621), according to manufacturer’s instructions. Gibson assembly fragments were generated by PCR using KAPA HiFi HotStart Ready Mix and relevant primers (Table S2) on plasmids containing the corresponding sequences. The IRE hairpin was added to one end of the plasmid backbone fragment via an extended PCR primer. After assembly of the parental plasmids, slicing sites were inserted as 24-bp sequences through site-directed mutagenesis by PCR (Table S2) to generate each of the eight resultant plasmid variants. Plasmid sequences were verified by Sanger sequencing.

Guide−passenger RNA duplexes for transfection were generated in the same way as those used for RISC assembly, but without any ^32^P-radiolabeled guide RNAs. To direct the correct strand selection by AGO2^87^ while minimizing variations in loading, identical mismatches were introduced in the duplex for the wildtype and mutant variants of each guide pair.

HeLa cells were plated at 30% confluency, and HEK293T cells were plated at 20% confluency, each in 96-well plates with 100 µL of media per well. Cells were transfected 24 h after plating, using Lipofectamine 2000, according to manufacturer’s instructions. Each well of cells was co-transfected using 0.18 µL of Lipofectamine 2000, 60 ng of reporter plasmid, and 0.6 pmol of carrier lsy-6 duplex, with or without 1.2 pmol of the RNA duplex being tested, mixed in Opti-MEM to a total volume of 50 µL. After 7 h of transfection, cells were treated for iron control. Stock solutions of 26.3 mg/mL (40 mM) desferroxamine mesylate (DFO; Sigma Aldrich, D9533) or 40 mg/mL ferric ammonium citrate (FAC; Sigma Aldrich, F5879) were prepared from powder, dissolved in 50% (v./v.) DMSO, and stored at −20°C. Cells were treated with 50 µL of either 3.3 µg (5 nmol) of DFO or 2.5 µg of FAC mixed in DMEM supplemented with 10% FBS, bringing the final total volume of media in each well to 200 µL. After 17 h of treatment, cells were examined and harvested for luciferase assays.

Fluorescence from sfGFP in cells transfected with context-A reporters was imaged using the EVOS M7000 Imaging System (Invitrogen). Fluorescence microscopy images were analyzed using ImageJ 1.53n, with the display intensity set to the range of 800−1500 out of a 15-bit range (0−32767) uniformly across all images.

Luciferase assays were conducted using the Dual-Luciferase Reporter assay system (Promega, E1960) according to manufacturer’s instructions. Cell lysates were transferred to a 96-well reading plate, and luminescence was measured using the GloMax Discover microplate reader (Promega) with the default parameters for dual-luciferase assays. Fifteen measurements of cell lysates without any expressed luciferase were used to estimate the background reading levels of firefly luciferase and NanoLuc luciferase, and the mean background for each was respectively subtracted from all experimental readings before analysis in R.

#### Analysis of RNA backbones in high-resolution structure models of AGO−RNA complexes

Structure models were downloaded from the RCSB Protein Data Bank. RNA backbone torsion angles were measured using the web 3DNA 2.0 server,^85^ followed by analysis in R. RNA backbone conformer analyses were conducted using MolProbity 4.2.^86^ Iterative-build composite electron density omit maps (2*mF*_o_ − *DF*_c_) were calculated in PHENIX 1.20,^81^ using the phenix.composite_omit_map command with no refinement, and rendered in PyMOL 2.4.1 at 1.6 *σ* for all models generated using crystallography.

#### Prediction of RNA secondary structure and base-pairing energy

Minimum free energy secondary structures and corresponding predicted standard free energy (Δ*G*°_37°C_) values for slicing target RNAs were predicted *in silico* using the RNAfold program in the ViennaRNA 2.0 package.^84^ Standard free energy of secondary structure involving the seed-match region was approximated by multiplying the total predicted standard free energy by the fraction of total base-pairs located within the seed-match region. Predicted Δ*G*°_37°C_ of sequence regions pairing to their complement were calculated using the RNAeval program in the ViennaRNA 2.0 package, corrected for the initiation and/or symmetry energy penalties (4.09 and 0.43 kcal mol^−1^, respectively^89^) when imposed.

## QUANTIFICATION AND STATISTICAL ANALYSIS

All radioactive gels and filters were imaged using Typhoon FLA 7000 (GE Healthcare), Typhoon FLA 9500 (GE Healthcare), or Amersham Typhoon (Cytiva) phosphorimager. Images were quantified using either the Multi Gauge (Fujifilm) or ImageQuant TL (Cytiva) software; for each band or filter, background signal was estimated as the mean signal of two equal-area regions drawn outside of the associated band or filter, and subtracted from the raw signal. All error bars represent 95% CIs calculated as 1.96 × standard error of the mean. Other specific statistical parameters, including the number of data points or replicates (*N*), statistical test used, and statistical significance (*p* value) are reported as appropriate in the corresponding figures and legends.

## SUPPLEMENTAL INFORMATION

**Table S1. Fitted parameters of slicing kinetics with different guides and targets, related to Figures 1, 2, 5 and 6**

**Table S2. Oligonucleotides used in this study, related to STAR Methods**

**Video S1. Conformational changes occurring during the transition from the two-helix state to the slicing-competent state, related to Figure 1**

## Notes

### Competing Interest Statement

D.P.B. has equity in Alnylam Pharmaceuticals, where he is a co-founder and advisor. P.Y.W. declares no competing interests.

### Summary of Updates

Introduction and discussion sections updated; other minor revisions.

