## Supplemental Figures and Legends for "The guide RNA sequence dictates the slicing kinetics and conformational dynamics of the Argonaute silencing complex"

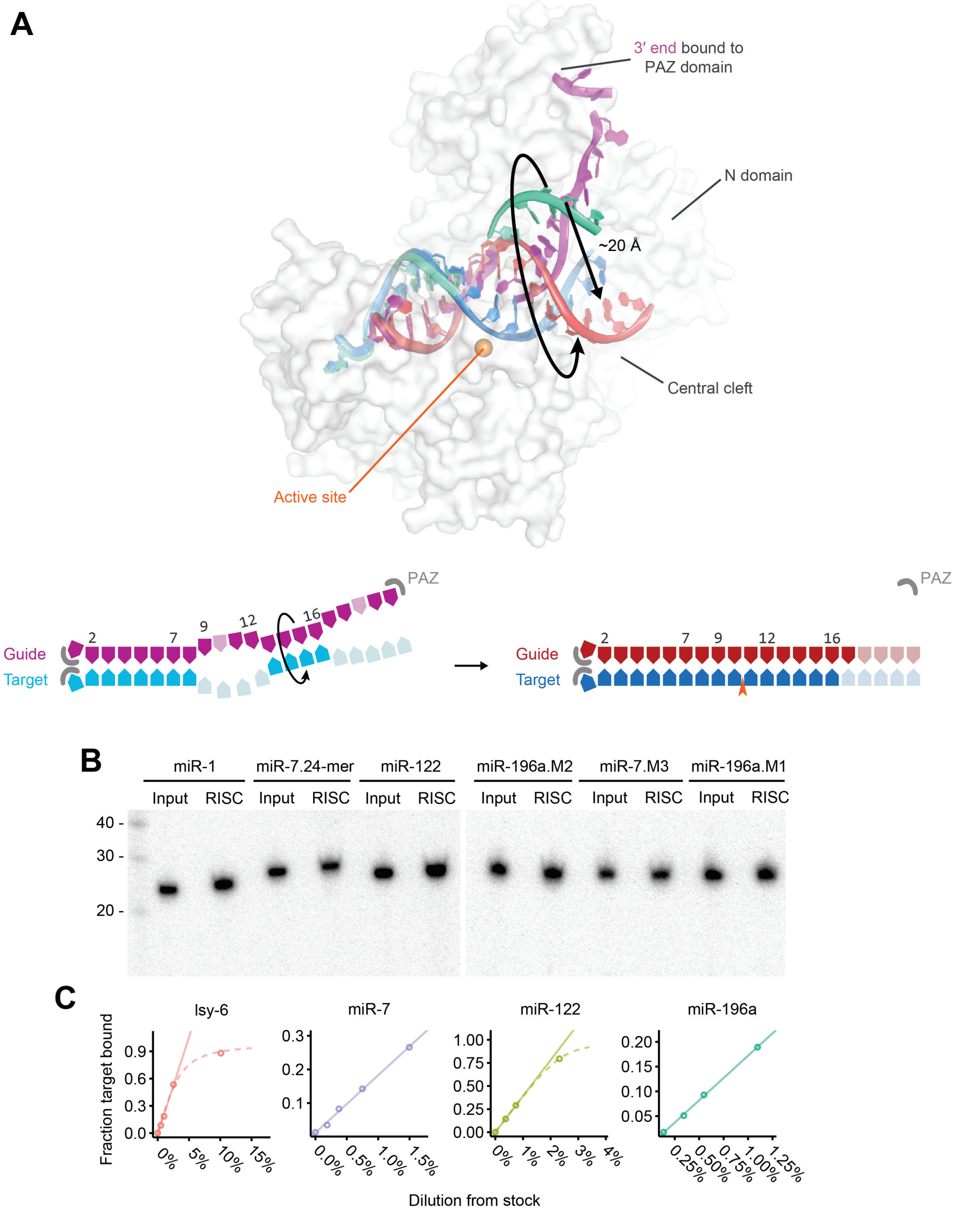


##### Figure S1. Structure and purification of RISCs, related to Figure 1

(A) Changes in RNA as RISC–target complex transitions from the two-helix state to the fully paired state. Structure model of RISC engaged with a slicing target in the two-helix state, prior to attaining the fully paired state (human AGO2, 6N4O [S1]), is show together with the RNA components of the centrally paired state (*A. thaliana* AGO10, 7SWF [S2]), which were superimposed by aligning at the MID and PIWI domains of the AGO. RNA base-pairing diagrams corresponding to the two states are on the right. AGO2 is shown as a light gray surface. The catalytic tetrad in the active site is shown as orange space-filling models in the structure, and as an orange caret in the pairing diagram. In the two-helix state, the guide is in magenta and the target in cyan; in the fully paired state, the guide is in red and the target in blue. Nucleotides not resolved or omitted in the structure models are shown in a lighter shade of color in the pairing diagrams on the right. Major movements of the second helix, required to assume the slicing-competent state, are indicated with black arrows.

(B) Examples of denaturing electrophoretic analysis of radiolabeled miRNAs, comparing for each miRNA the mobility and quantity of miRNA loaded into RISC with the mobility and quantity of miRNA within the corresponding input miRNA−passenger duplexes. Samples were run on an 8M urea, 15% polyacrylamide gel, and radiolabeled miRNA was visualized by phosphorimaging. Intensities of bands from the RISCs were compared to those of the known duplex concentrations to generate initial estimates of RISC concentrations.

(C) Representative results from filter-binding experiments used to quantify RISC concentration by titrating 1.0 nM end-labeled, seed-matched target. Data points that did not approach saturation (≤ 60% bound) were fit by linear regression (solid lines). Datasets with saturation behavior were also fit to a quadratic equation by nonlinear least-squares regression (dashed lines).

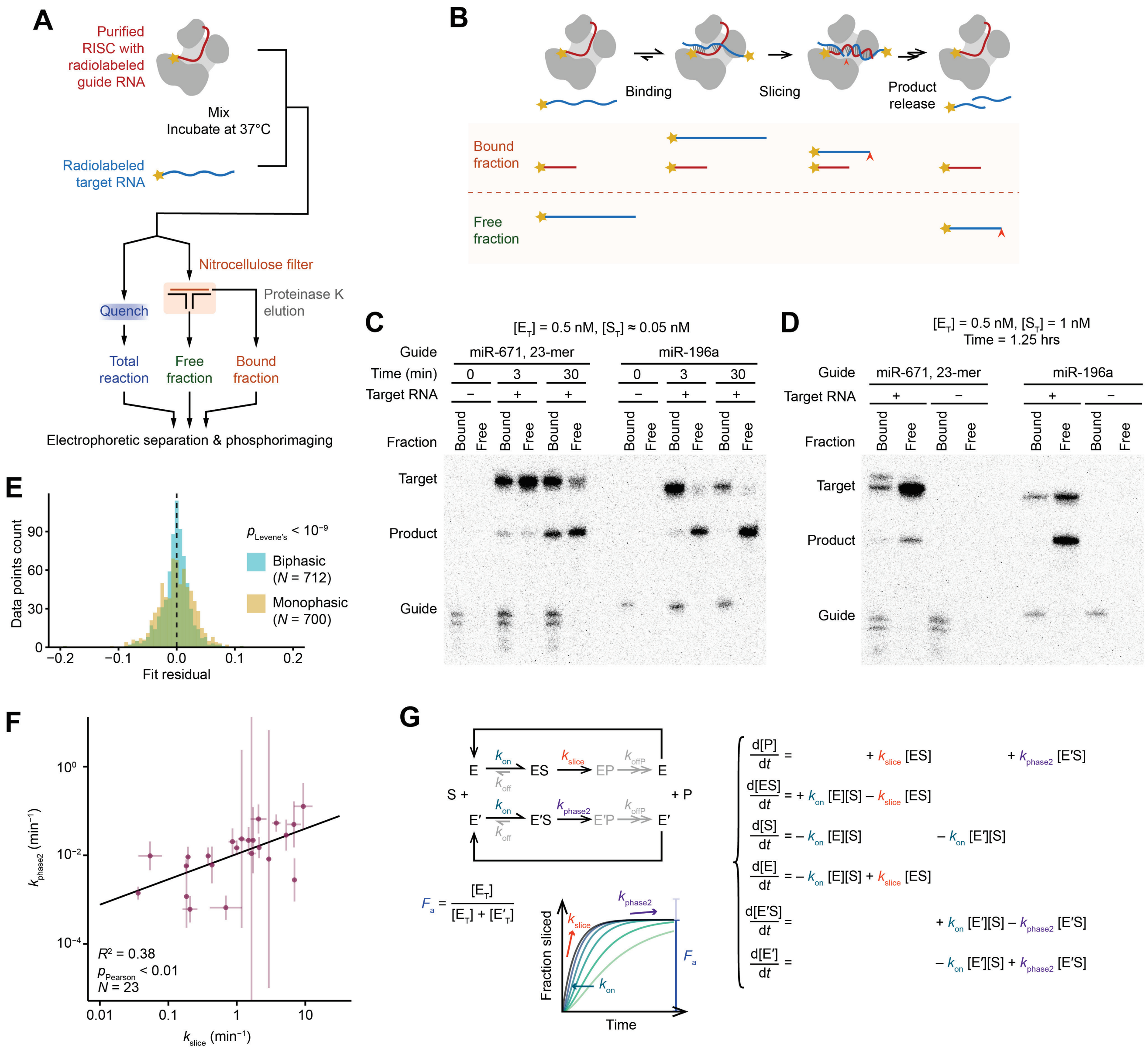

**Figure S2. Validation of ODE models, related to Figure 1**

(A) Schematic of filter-binding experiments of slicing assays; otherwise as in Figure 1B.

(B) Schematic of expected locations following nitrocellulose filtration of RNAs of each species along the slicing reaction. Radiolabeled sliced product is indicated with an orange caret at the cleavage site; otherwise, as in Figure 1B.

(C) Filter-binding results of single-turnover slicing assays. miR-671 (23-mer) had slow binding kinetics in slicing assays, whereas miR-196a had faster binding kinetics (Figure S4). At time 0, no target had been added.

(D) Filter-binding results of slicing assays in which the target substrate was in excess over RISC; otherwise, as in (C). None of the miRNAs underwent substantial unloading from AGO2 upon binding slicing targets [S3].

(E) Analysis of the improved fit observed when fitting slicing results to a biphasic, rather than a monophasic, ODE model. For each model, the distribution of fit residuals for time points at least three times the value of $\tau_{slice}$ is plotted. $N$ indicates the numbers of data points included. The $p$ value was calculated using Levene's test for equality of variances.

(F) Relationship between second-phase slicing kinetics and the first-phase kinetics. Error bars indicate 95% CIs of model-fitting. Datasets with insufficient data to fit for $k_{phase2}$ are excluded. The Pearson correlation and linear regression results are plotted.

(G) Full ODE model used to fit single-turnover slicing assay datasets, incorporating biphasic kinetics. A second, slow phase is modeled as a small fraction (1 – $F_a$) of defective enzyme (E'); otherwise, as in Figure 1E. Values for the elemental rate constant of this second phase ($k_{phase2}$ or kph2 values) are reported (Table S1).

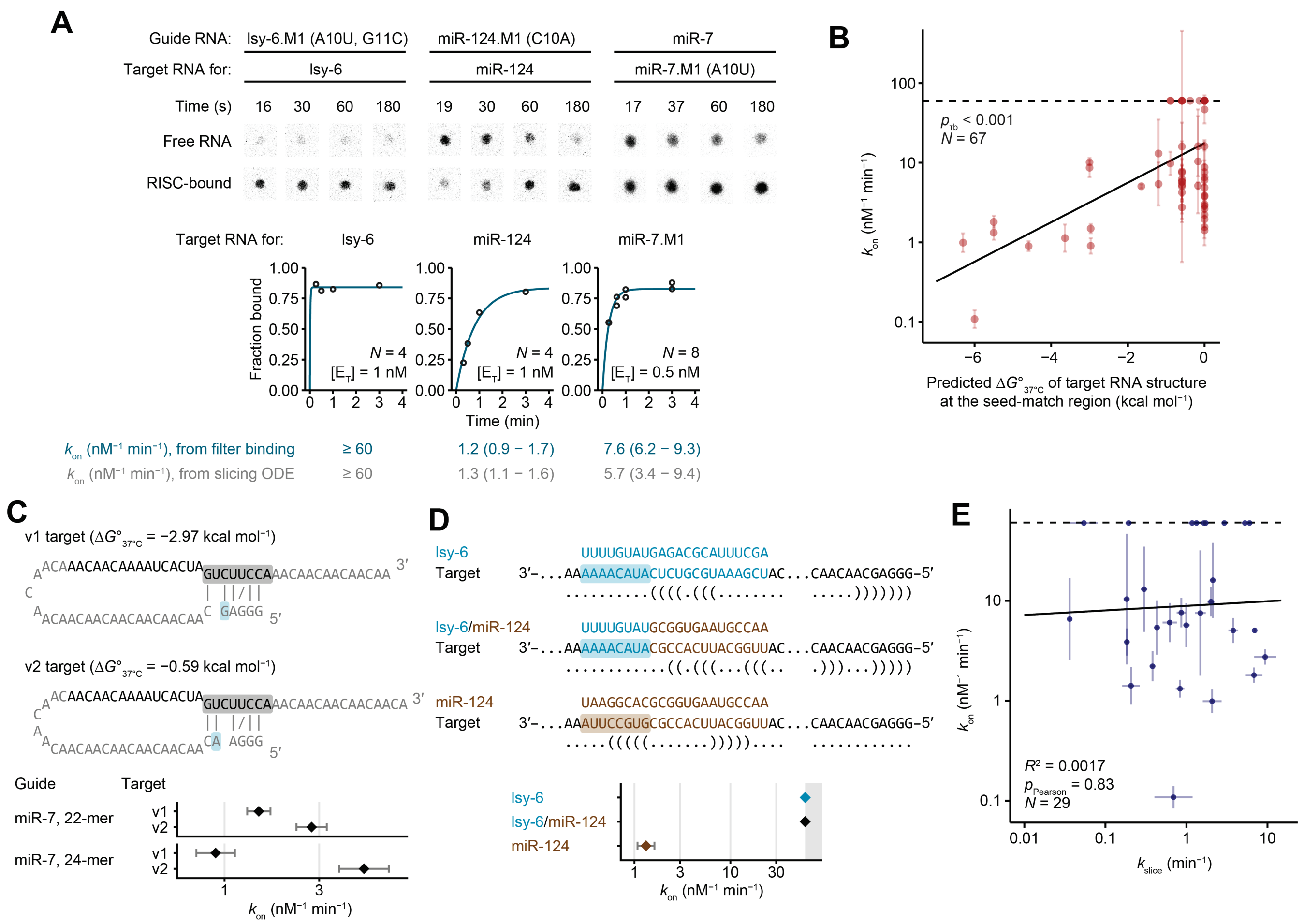


#### Figure S3. Influence of target secondary structure on association kinetics, related to Figure 1

(A) Orthogonal confirmation of $k_{on}$ values from ODE model fitting by filter binding. Shown are representative filter-binding results and their corresponding kinetic binding plots. Data points were fit to an exponential equation. Fit results of $k_{on}$ are shown together with the values determined from ODE model fitting. Error ranges indicate 95% CIs of model-fitting. Association kinetics of lsy-6 were too fast to be resolved by either our ODE model fitting or our filter-binding assays, and are therefore shown as and plotted at the diffusion limit. The number of data points ($N$) and RISC concentrations are shown. For each assay, the concentration of target RNA was 10-fold below the RISC concentration.

(B) The relationship between the $k_{on}$ values measured from ODE model fitting and the predicted standard free energy of secondary structure ($\Delta G°_{37°C}$) at the seed-matched site in the target RNA. The linear best-fit line is plotted, and the $p$ value of the non-parametric Kendall's tau-b test is shown, which accommodates the non-normal distribution and the existence of ties in the data. The number of data points is indicated as $N$. The diffusion limit for $k_{on}$ is shown as a dashed line.

(C) Influence of a single target-RNA substitution that weakens predicted secondary structure at the seed-match region. Two versions of the slicing target for miR-7 are shown, each with its predicted secondary structure and associated standard free energy. For each target, the perfectly complementary target site is colored in black, and the seed-match region is highlighted in gray. The single-nucleotide substitution predicted to reduce stability of the secondary structure at the seed-match region is highlighted in blue. The $k_{on}$ values for each target, as measured from ODE model fitting, are shown, similarly to Figure 2C.

(D) Results from a chimeric guide isolating the effect of pairing to the seed region on association kinetics. The miRNA and target sequences for lsy-6 (turquoise), miR-124 (brown), and their chimera are shown. The seed-match region is highlighted for each target. Predicted secondary structure at the relevant regions of each target RNA is indicated in dot-bracket notation.

(E) Insignificant correlation observed between $k_{slice}$ and $k_{on}$ values for perfectly complementary targets; otherwise as in (B) and Figure S2F.

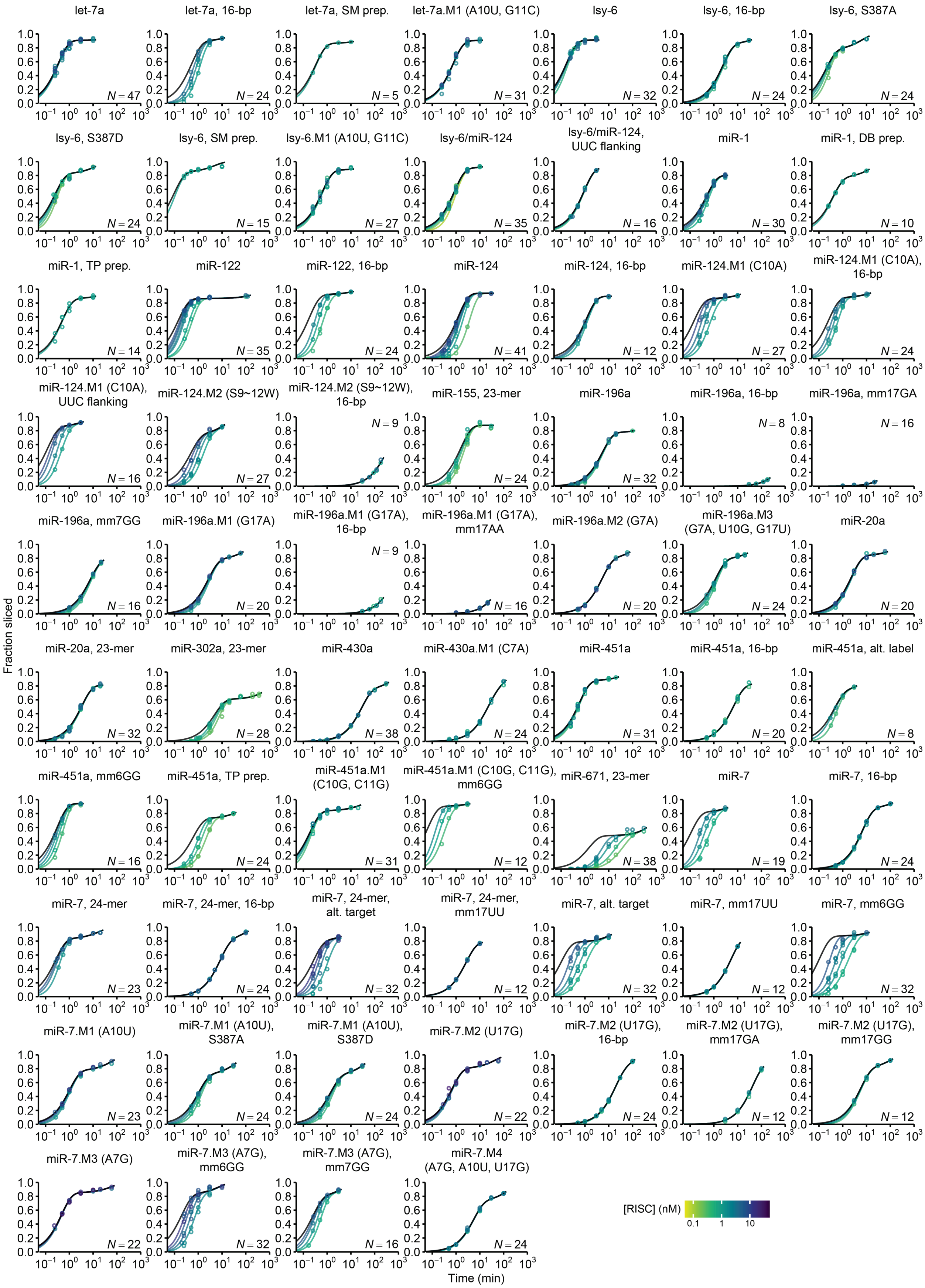
let-7a
N = 47
let-7a, 16-bp
N = 24
let-7a, SM prep.
N = 5
let-7a.M1 (A10U, G11C)
N = 31
lsy-6
N = 32
lsy-6, 16-bp
N = 24
lsy-6, S387A
N = 24
lsy-6, S387D
N = 24
lsy-6, SM prep.
N = 15
lsy-6.M1 (A10U, G11C)
N = 27
lsy-6/miR-124
N = 35
lsy-6/miR-124, UUC flanking
N = 16
miR-1
N = 30
miR-1, DB prep.
N = 10
miR-1, TP prep.
N = 14
miR-122
N = 35
miR-122, 16-bp
N = 24
miR-124
N = 41
miR-124, 16-bp
N = 12
miR-124.M1 (C10A)
N = 27
miR-124.M1 (C10A), 16-bp
N = 24
miR-124.M1 (C10A), UUC flanking
N = 16
miR-124.M2 (S9~12W)
N = 27
miR-124.M2 (S9~12W), 16-bp
N = 9
miR-155, 23-mer
N = 24
miR-196a
N = 32
miR-196a, 16-bp
N = 8
miR-196a, mm17GA
N = 16
miR-196a, mm7GG
N = 16
miR-196a.M1 (G17A)
N = 20
miR-196a.M1 (G17A), 16-bp
N = 9
miR-196a.M1 (G17A), mm17AA
N = 16
miR-196a.M2 (G7A)
N = 20
miR-196a.M3 (G7A, U10G, G17U)
N = 24
miR-20a
N = 20
miR-20a, 23-mer
N = 32
miR-302a, 23-mer
N = 28
miR-430a
N = 38
miR-430a.M1 (C7A)
N = 24
miR-451a
N = 31
miR-451a, 16-bp
N = 20
miR-451a, alt. label
N = 8
miR-451a, mm6GG
N = 16
miR-451a, TP prep.
N = 24
miR-451a.M1 (C10G, C11G)
N = 31
miR-451a.M1 (C10G, C11G), mm6GG
N = 12
miR-671, 23-mer
N = 38
miR-7
N = 19
miR-7, 16-bp
N = 24
miR-7, 24-mer
N = 23
miR-7, 24-mer, 16-bp
N = 24
miR-7, 24-mer, alt. target
N = 32
miR-7, 24-mer, mm17UU
N = 12
miR-7, alt. target
N = 32
miR-7, mm17UU
N = 12
miR-7, mm6GG
N = 32
miR-7.M1 (A10U)
N = 23
miR-7.M1 (A10U), S387A
N = 24
miR-7.M1 (A10U), S387D
N = 24
miR-7.M2 (U17G)
N = 22
miR-7.M2 (U17G), 16-bp
N = 24
miR-7.M2 (U17G), mm17GA
N = 12
miR-7.M2 (U17G), mm17GG
N = 12
miR-7.M3 (A7G)
N = 22
miR-7.M3 (A7G), mm6GG
N = 32
miR-7.M3 (A7G), mm7GG
N = 16
miR-7.M4 (A7G, A10U, U17G)
N = 24
Fraction sliced
Time (min)
[RISC] (nM)
0.1
1
10

**Figure S4. Results of slicing assays for all sequences and conditions tested, related to Figure 1**

Colors and details are as in Figure 1D, but in log-scale for the x-axes. All data points are shown. Values of parameters obtained from model fitting are provided (Table S1). SM, DB, and TP prep indicate preparations from Sean McGeary, Daniel Briskin, and Thy Pham, respectively. Alt. label indicates radiolabeling as either a 5′ monophosphate or an $m^7G$ cap. Alt. target indicates different target flanking sequence designs. Results of miR-7, miR-451a, miR-124, and miR-196a RISCs are replotted from Figure 1D in log-scale.

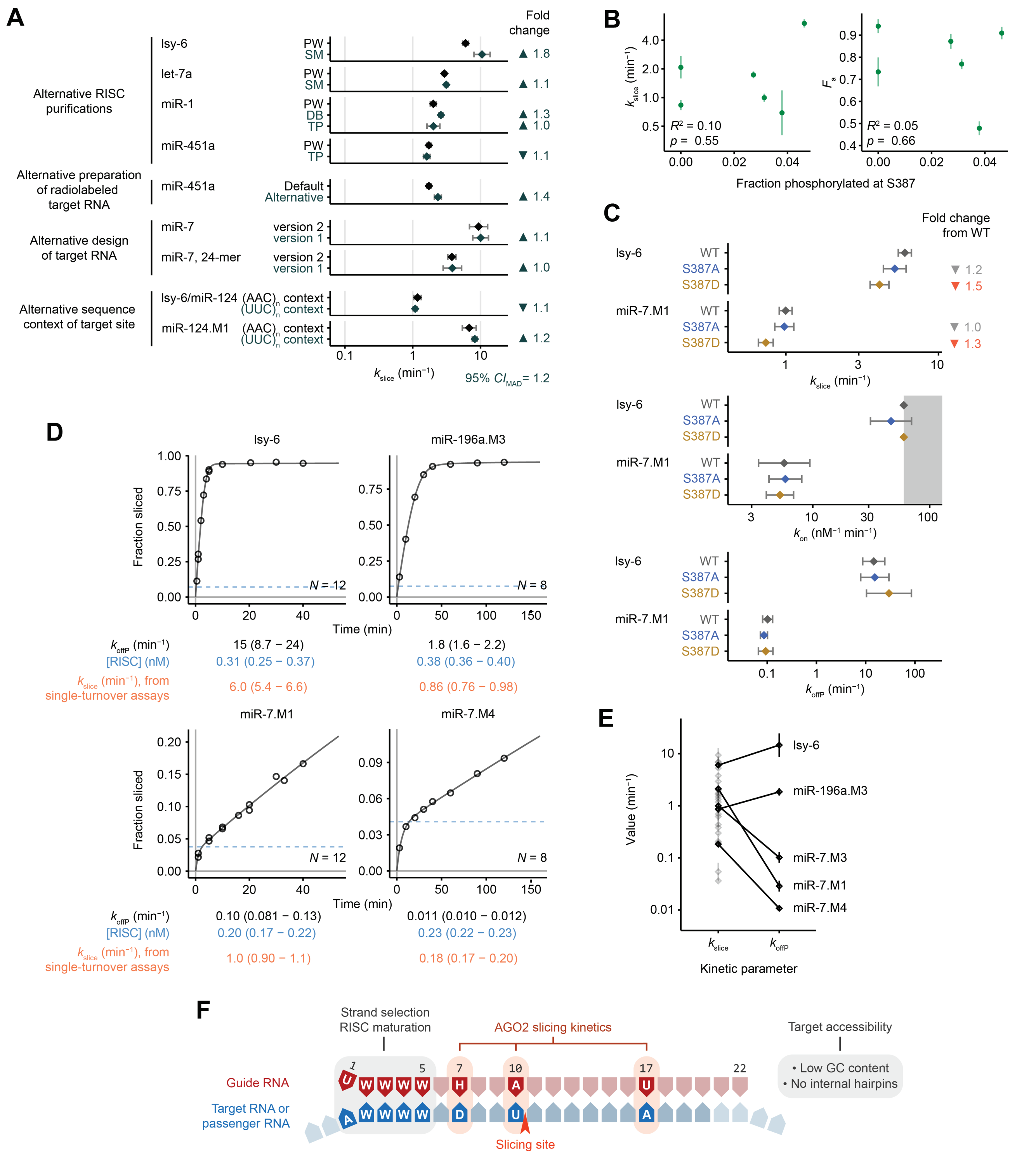

### Figure S5. Relationship between $k_{slice}$ with other factors or kinetic parameters, related to Figure 2

(A) Reproducibility of $k_{slice}$ measurements. For the alternative RISC purifications, the initials indicate the person who purified the RISC (PW, Peter Wang; SM, Sean McGeary; DB, Daniel Briskin; TP, Thy Pham). The summarized 95% CI of background variation based on the median absolute deviation (MAD) is shown. Otherwise, this panel is as in Figure 2C. Values in black are replotted from Figure 2D.

(B) Evidence that the fraction of phosphorylated S387 is too low to perceptively affect slicing. Plots show insignificant correlation between either $k_{slice}$ or $F_a$ values and the fraction of phosphorylated S387, as measured by mass spectrometry of six purified RISCs. Error bars indicate 95% CIs of model-fitting. Coefficients of determination ($R^2$ values) derived from linear regression are shown, along with the $p$ values determined using the $t$-test.

(C) Fold changes in kinetic parameters observed upon either the phospho-null (S387A; blue) or the phosphomimetic (S387E; yellow) substitution of AGO2 S387. The diffusion limit for $k_{on}$ is shown as gray shading; otherwise, as in Figure 2C. $k_{slice}$ values with wildtype AGO2 are replotted from Figure 2D.

(D) Multiple-turnover slicing assays for four different guides. Black lines indicate results of fitting to a multiple-turnover ODE model, using $k_{slice}$ and $k_{on}$ values previously fit from single-turnover assays. For each dataset, the number of data points is indicated as $N$. Results from ODE model fitting of $k_{offP}$ and RISC concentrations are shown with 95% CIs, alongside the previously determined $k_{slice}$ values. For each reaction, the initial concentration of target RNA was 4 nM; the fraction of target RNA corresponding to the RISC concentration, expected to react during the first catalytic turnover, is indicated with a blue dashed line.

(E) Comparison of values measured for $k_{slice}$ and $k_{offP}$. Error bars indicate 95% CIs of model-fitting. Values for the five guides with measured $k_{offP}$ values are plotted in black, whereas those of other guides are in gray.

(F) siRNA sequence features associated with more effective RNAi in cultured cells [S4]. Features previously mechanistically explained are highlighted in gray, and those newly explained in this work highlighted in orange. Otherwise, this panel is as in Figure 2E.

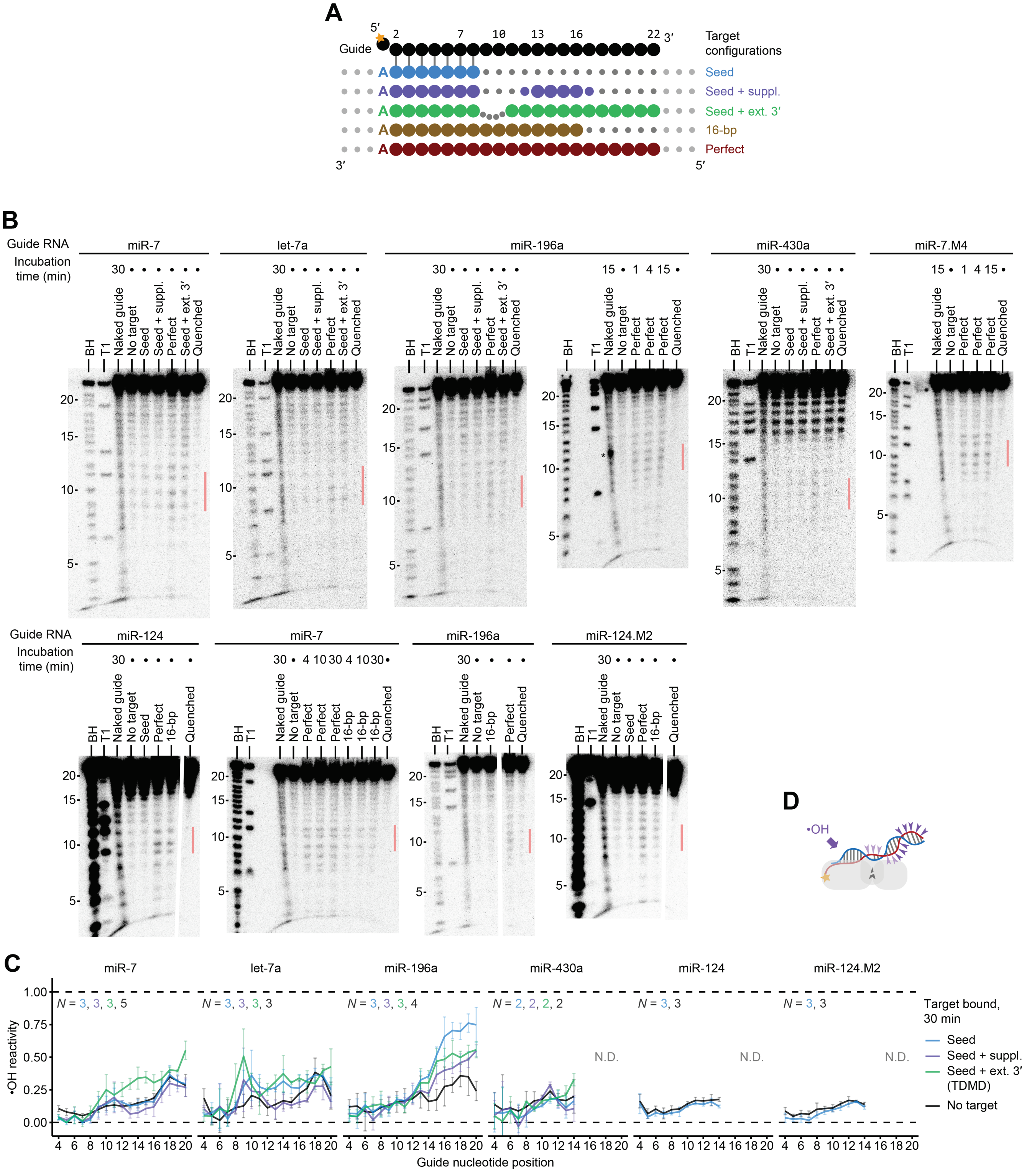


#### Figure S6. Hydroxyl-radical footprinting used to probe RISC conformations, related to Figures 4 and 6

(A) Pairing configurations for types of targets tested in footprinting experiments.

(B) Representative results from hydroxyl-radical footprinting experiments. Positions 9−11 are highlighted with orange bars on the right. Ladders from partial base-hydrolysis (BH) or partial RNase T1-digest (T1) are shown. Time of incubation with the specified target before footprinting is shown for each condition (with a dot indicating times identical to those of the preceding lanes). A non-specific degradation band in one replicate of the naked miR-196a sample preparation is indicated with an asterisk. Lanes corresponding to samples not relevant to this work are removed where indicated with space.

(C) Reactivity values observed after AGO2$^{D669A}$−miRNA was engaged with targets with seed-only, seed+supplementary, or seed and extensive 3′ complementarity. No-target values are replotted from Figure 4D and 6F. Colors are as in (B). Otherwise, this panel is as in Figure 4D.

(D) Schematic of reactivity patterns expected with seed and extensive 3′ pairing based on crystallographic structure models [S5], similar to Figure 4C.

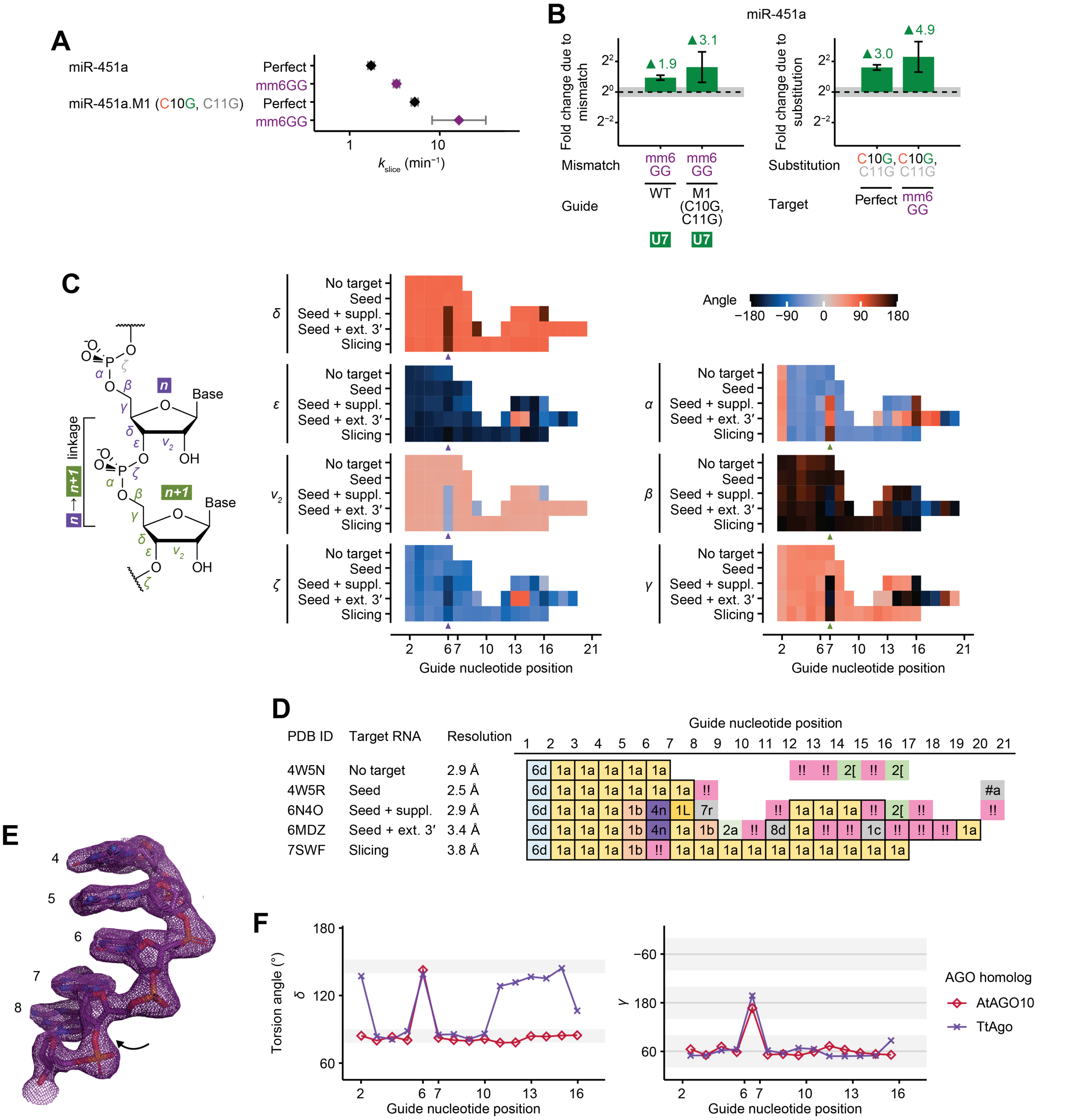


**Figure S7. Additional evidence that enhancement of slicing by weak pairs or mismatches at positions 6−7 aligns with local backbone distortion, related to Figure 5**

(A) $k_{slice}$ values of perfect targets or targets with a position-6 G:G mismatch, comparing results for miR-451a with those for its mutant with substitutions at positions 10 and 11. Otherwise, this panel is as in Figure 5A. Values with perfectly complementary targets are replotted from Figure 2D.

(B) Fold-change comparisons between $k_{slice}$ values in (A); otherwise, as in Figure 5B.

(C) RNA backbone torsion angles at guide positions within the five structure models analyzed in Figure 5C. Values are only shown for nucleotides engaged in base-pairs or, in the case of the no-target state, positions 2−7. Values at the linkage between positions 6−7 are indicated with a small triangle below.

(D) Conformer assignments at guide-RNA linkages within the five structure models analyzed in Figure 5C, based on the two-character nomenclature by the RNA Ontology Consortium; for example, 1a corresponds to standard A-form, whereas !! corresponds to outliers not found in the standard set of consensus conformers [S6,S7]. Positions engaged in pairing are framed with black squares.

(E) Similar backbone distortion of the DNA guide as *Thermus thermophilus* Ago assumes the slicing-competent conformation. Structure model of the guide backbone at positions 4−8 is shown with electron density omit maps (4NCB, at 2.2 Å resolution) [S8], similarly to Figure 5D. Movement to form the kink is shown with a black arrow.

(F) *δ* and *γ* torsion angles observed in the guide backbone for *Thermus thermophilus* Ago in the slicing-competent conformation (4NCB, purple), compared to those replotted for *Arabidopsis thaliana* AGO10 (7SWF, deep pink) from Figure 5C. Otherwise, this panel is as in Figure 5C.
